# Profiling subcellular localization of nuclear-encoded mitochondrial gene products in zebrafish

**DOI:** 10.1101/2021.12.22.473872

**Authors:** Barbara Uszczynska-Ratajczak, Sreedevi Sugunan, Monika Kwiatkowska, Maciej Migdal, Silvia Carbonell-Sala, Anna Sokol, Cecilia L. Winata, Agnieszka Chacinska

**Author notes:** These authors contributed equally (Barbara Uszczynska-Ratajczak, Sreedevi Sugunan).

## Abstract

Most mitochondrial proteins are encoded by nuclear genes, synthetized in the cytosol and targeted into the organelle. The import of some, but not all, nuclear-encoded mitochondrial proteins begins with translation of messenger RNAs (mRNAs) on the surface of mitochondria. To characterize the spatial organization of mitochondrial gene products in zebrafish (*Danio rerio*), we sequenced RNA from different cellular fractions. Our results confirmed the presence of nuclear-encoded mRNAs in the mitochondrial fraction, which in unperturbed conditions, are mainly transcripts encoding large proteins with specific properties, like transmembrane domains. To further explore the principles of mitochondrial protein compartmentalization in zebrafish, we quantified the transcriptomic changes for each subcellular fraction triggered by the *chchd4a^-^/^-^* mutation, causing the disorders in the mitochondrial protein import. Our results indicate that the proteostatic stress further restricts the population of transcripts on the mitochondrial surface, allowing only the largest and the most evolutionary conserved proteins to be synthetized there. We also show that many nuclear-encoded mitochondrial transcripts translated by the cytosolic ribosomes stay resistant to the global translation shutdown. Thus, vertebrates, in contrast to yeast, are not likely to employ localized translation to facilitate synthesis of mitochondrial proteins under proteostatic stress conditions.

## Introduction

Despite the presence of mitochondrial DNA, the vast majority of mitochondrial proteins (∼99%) are encoded by the nuclear genome. Therefore, the biogenesis of functional mitochondria relies on the synthesis and import of those proteins, in their precursor form, into the organelle. Efficient protein import through a highly complex mitochondrial structure requires a system regulating transport and assembly of precursor proteins^1–3^. Following the synthesis on cytosolic ribosomes, precursor proteins are imported through the translocase of the membrane (TOM) complex, which is a common entry gate for all mitochondria-destined precursor proteins. After crossing TOM, precursor proteins are directed to specific mitochondrial compartments including outer mitochondrial membrane (OMM), inner mitochondrial membrane (IMM), intermembrane space (IMS) and matrix^4–6^.

Although the mitochondrial protein import has been extensively studied for decades, the cytosolic stage of this process is not fully understood^7^. At the same time, an efficient trafficking of RNA into specific compartments is a fundamental task of eukaryotic cells, as a precise distribution of mRNAs across organelles in the cell allows to synthetize proteins exactly, where they are needed^8^. As a consequence, numerous studies have indicated the presence of mRNAs encoding mitochondrial proteins, either on the mitochondrial surface or in its close proximity^9–13^. Moreover, mitochondria associated mRNAs have been proven to be active templates for protein synthesis^14^, whereas the cytosolic ribosomes observed at OMM^15,16^ are able to interact with the TOM complex^17^. Thus, in living cells the translation of nuclear-encoded proteins can occur at the surface of mitochondria and might be directly coupled to their import into this organelle^14,18^. Interestingly, it seems that the physicochemical properties of mitochondrial proteins, including hydrophobicity or folding speed, might determine the protein import route^19–22^. Despite these findings, mitochondrial protein import remains almost completely unexplored for higher eukaryotes, as this process was mainly studied using yeast^9–14,20,23–25^ and mammalian cell^26–28^ models.

Biogenesis of mitochondria is a precisely controlled process that largely depends on a fine-tuned balance between synthesis and degradation of many cellular proteins^29^. Disorders in the mitochondrial protein import can trigger serious consequences for the cell, including energetic deficiencies and proteostatic stress response induced by the accumulation of precursor proteins in the cytosol^30^. Although there are various cellular mechanisms that can reduce the accumulation of mis-targeted mitochondrial proteins^30–32^, we still do not understand the impact of mitochondrial stress on the subcellular localization of mRNAs encoding mitochondrial proteins. In general, the knowledge on how a cell maintains the biogenesis of mitochondria under the proteostatic stress is very limited. At the same time, various alternative mitochondrial protein targeting routes that protect and redirect mis-targeted proteins outside mitochondria exist^33^. Surprisingly, many of them involve interaction with endoplasmic reticulum (ER)^34,35^, in particular under proteostatic stress conditions^36–38^.

Here, we studied the subcellular localization of nuclear-encoded mitochondrial mRNAs in zebrafish to uncover transcripts that are enriched and thus, are likely to be translated on the mitochondrial surface. Additionally, this study for the first time reports the dynamics of subcellular localization of mitochondrial transcripts in physiological conditions and under protein import deficiency. To assess the impact of mitochondrial protein import disorders on the mitochondria-associated transcripts, we took advantage of *chchd4a^-^/^-^* zebrafish mutant line^39^. As previously described, loss of function of the *chchd4a* paralogue affects the activity of the MIA pathway – an essential route for import and assembly of mitochondrial proteins in the IMS. Here, we further explore the transcriptomic response triggered by *chchd4a^-^/^-^* mutation within the context of subcellular mRNA localization. Taken together, this study broadens our knowledge about the principles and dynamics of distributing mRNAs encoding mitochondrial proteins across the cell in vertebrates.

## Materials and Methods

### Zebrafish lines and maintenance

Zebrafish wild type line from AB background and *chchd4a^+/−^* mutant line^39^ in the *Tg(Xla*.*Eef1a1*:*mlsEGFP)cms1* transgenic background^40,41^ were maintained in the IIMCB zebrafish core facility (License no. PL14656251) in accordance with institutional and national ethical and animal welfare guidelines. The optimization of biochemical fractionation was carried out using the AB line. 5 days post fertilization (dpf) larvae obtained from an in-cross of *chchd4a^+/+^* siblings in homozygous *Tg(Xla*.*Eef1a1*:*mlsEGFP)cms1* background were used as the WT reference for fractionation, RNA sequencing and RT-qPCR validations. Embryos obtained from the in-cross of heterozygous *chchd4a^+/−^* in homozygous *Tg(Xla*.*Eef1a1*:*mlsEGFP)cms1* background were observed under a fluorescent microscope at 4 dpf to select *chchd4a*^-/-^ individuals based on the characteristic GFP-containing abnormal mitochondrial structures described previously^39^. The larvae of *chchd4a^+/+^* and *chchd4a^-/-^* in *Tg(Xla*.*Eef1a1*:*mlsEGFP)cms1* background were grown in an incubator at 28°C until 5 dpf and used for fractionation-sequencing.

### Subcellular fractionation of 5dpf larvae and isolation of membrane-bound and high-speed fractions

All steps were carried out at 4°C unless otherwise specified. For subcellular fractionation, around 250 zebrafish larvae at 5 dpf were anesthetized using Tricaine (Sigma A5040) in E3 medium, washed thrice with Ringer’s solution (116 mM NaCl, 2.9 mM KCl, 5.0 mM HEPES, pH 7.6) and incubated in cold isolation buffer (IB) composed of 220 mM mannitol, 70 mM sucrose, 20 mM HEPES (pH 7.6), for at least 30 min. The larvae were suspended in 4ml of the isolation buffer with EDTA (IB-E; 20mM HEPES (pH 7.6), 220mM mannitol, 70mM sucrose, 1mM EDTA, supplemented freshly with 2mM PMSF) or in isolation buffer with magnesium (IB-M; 20mM HEPES (pH 7.6), 220 mM mannitol, 70mM sucrose, 5 mM MgCl_2,_ supplemented freshly with 2 mM PMSF, 200µg/ml cycloheximide (CHX) and 10U/ml RiboLock RNase inhibitor). Then the larvae were homogenized in a 5 ml Dounce glass homogenizer equipped with a tight-fitting glass pestle by giving 15 strokes. The homogenate (total) was centrifuged twice at 1500 X g for 10 min to remove nuclei and debris to obtain the post-nuclear supernatant (PNS). The PNS was further centrifuged at 14,000 X g for 15 min to pellet the membrane-bound (MB) fraction that is expected to contain mitochondria. The resulting supernatant was centrifuged at 100,000 X g for 60 min to obtain the second pellet – the high-speed (HS) and the cytosol as supernatant.

Isolation of crude and EDTA-stripped MB and HS fractions from WT and *chchd4a^-/-^* were performed as described above, with the following modification. The IB-M and IB-E buffers were additionally supplemented with 2 mg/ml of BSA.

Further, obtained subcellular fractions were used for protein or RNA isolation. For protein steady-state level analysis, the concentration of the total, cytosolic, and solubilized in IB buffer MB and HS fractions were estimated by Bradford assay or Direct Detect spectrophotometer (Millipore). Protein samples were solubilized in Laemmli buffer with 50 mM DTT, denatured at 65°C for 15 min, and subjected to SDS–PAGE and Western blotting. RNA isolation procedure is described below.

### RNA isolation

RNA was isolated from subcellular fractions using the TRI Reagent solution (ThermoFisher Scientific, AM9738). 500 µl of TRI Reagent was mixed with the sample and kept at room temperature for 8 min. into each sample, 100µl of chloroform was added, mixed and shaken for 15 seconds, followed by incubation at room temperature for 10 min. The samples were then centrifuged at 12,000 X g for 15 min at 4°C. Upon transfer of the aqueous phase to a new tube, 250 µl of isopropanol was added, followed by immediate vortexing for 15 sec. The samples were precipitated in −20°C overnight. The next day, samples were centrifuged at 20,000 X g for 15 min at 4 °C. After discarding the supernatant, the pellets were washed with 500 µl of 75 % ethanol. The RNA pellets were subsequently dissolved in an appropriate volume of RNase free water and treated with RNase-Free DNase (Qiagen, 79254) by incubation at room temperature for 10 min to avoid genomic DNA contamination. Further, the RNA samples were cleaned and concentrated using Agencourt RNAClean XP beads (Beckman Coulter, A63987) following the 1.8 X reaction volume protocol. The RNA concentration was estimated by Quantus Fluorometer (Promega) or Nanodrop Spectrophotometer (ThermoFisher Scientific). For library generation, the integrity of RNA was verified by 2200 Tapestation system (Agilent Technologies). The RNA samples were stored in −80°C until further use.

### RT-qPCR

cDNA synthesis was done using Verso cDNA synthesis kit (ThermoFisher Scientific, AB-1453/B) with an input of 250 ng of total RNA. ERCC Spike-In (Thermofisher Scientific, 4456739) controls (0.5μl of mix 1 or mix 2 from 1:100 dilution) were added into RNA samples. Random hexamers were used as primers for reverse transcription. Gene-specific primers for qPCR were designed with the help of primer BLAST software (Table S1). qPCR was performed using PowerUp SYBR Green Master Mix (ThermoFisher Scientific, A25742). The reaction was carried out in the 7900HT Real-Time PCR Instrument (Applied Biosystems) using default standard cycling mode conditions. Relative enrichment of RNA was calculated by normalizing Ct values against endogenous control using the 2^-ΔΔCt^ method^42^. For mitochondria versus MB comparison, *mt-atp8* was used as reference gene. While for RNA-seq validations, an ERCC Spike-In transcript (ERCC-00096) was used as reference.

### Short-read RNA sequencing

As input, 2 µg of RNA from each sample was incorporated with ERCC ExFold RNA Spike-In mixes (Thermofisher Scientific, 4456739) following the manufacturer’s guideline. The libraries were prepared using the KAPA Stranded mRNA-Seq kit (Kapa biosystems, KK8420) as follows. After polyA capture and RNA fragmentation step (6 min at 94°C), the first-strand cDNA was synthesized with random primers. Further, second-strand synthesis was carried out to convert the cDNA:RNA hybrid to double-stranded cDNA along with marking the second strand with dUTP. The samples were then cleaned up using 1.8 X Ampure bead-based protocol and immediately proceeded to A-Tailing to add dAMP to 3’-end of double-stranded cDNA fragments. Following A-Tailing, KAPA single indexed adaptor set A (KAPA biosystems, KK8701) was used for adaptor ligation. Samples were cleaned up twice using Ampure beads (1X bead-based). Next, the adaptor-ligated library DNA was amplified by 11 cycles of PCR. Finally, the library was cleaned up using Ampure beads (1X bead-based). Library fragment size distribution was confirmed by electrophoresis in Tapestation (Agilent Technologies). Library concentration was determined by both Quantus and qPCR (KAPA Library Quantification Kit Illumina® Platforms, KR0405). Sequencing was performed on the NextSeq 550 instrument (Illumina) using the v2 chemistry, resulting in an average of ∼100 M reads per library with 2 x 75 bp paired-end setup.

### RNA-seq data analysis

Raw Illumina NextSeq 550 reads were assessed for quality, adapter content and duplication rates with FastQC^43^. Next, raw Illumina reads were aligned to the reference transcriptome obtained using the masked DanRer11 (GRCz11) genome (plus the sequences of 96 ERCC Spike-In Controls) and Ensembl zebrafish gene annotation (v.101). The transcriptome sequence was prepared using the in house developed script for retrieving spliced transcript sequences. Whereas, the short-read mapping and quantification at the gene and isoform level was performed using Salmon^44^ (v.1.4.0) with default parameters. For the analysis of genome coverage short-reads were aligned to the masked DanRer11 (GRCz11) genome assembly using STAR^45^ (v.2.5.1) compiled for short-reads with following non-default parameters: -- outFilterMismatchNoverLmax 0.04 --alignIntronMin 20 --alignIntronMax 1000000 -- alignMatesGapMax 1000000 --outSAMunmapped Within --runThreadN 6. Illumina RNA-seq mapping statistics are summarized in supplementary Figure S1A and S1B. The genome coverage analysis (Figure S1C and S1D) was done using *bamstats* tool developed by Roderic Guigo’s group^46^.

Gene enrichment analysis was performed using DESeq2^47^ (version 1.30) and by directly comparing MB to HS fractions for wild type and *chchd4a^-/-^* samples, respectively. Only genes with total number of counts >50 across MB and HS samples were included in this analysis. Moreover, genes with a minimum fold change of log2 <=> +/− 1 and a maximum Benjamini-Hochberg corrected P-value of 0.05 were deemed to be classified as (highly) enriched in each fraction (Table S2). To control the quality of the analysis, we also distinguished moderately enriched genes with a minimum fold change of log2 <=> +/− (0.5-1) and a maximum Benjamini-Hochberg corrected P-value of 0.05. The analysis on the transcript level was performed using dominant transcripts – transcripts showing the highest expression levels across all the isoforms for a given gene. Raw Illumina RNA-seq data were deposited to GEO repository with the accession number GSE167587.

Previously generated RNA-seq data from whole unfractionated *chchd4a^-/-^* and wild type samples were quantified at the gene level using Salmon^44^ (v.1.4.0) with default parameters. Differentially expressed genes were identified using DESeq2^47^ (version 1.30). Only genes with a minimum fold change of log2 <=> 0.9, a maximum Benjamini-Hochberg corrected P-value of 0.05, and a minimum combined mean of 5 reads were deemed to be significantly differentially expressed (Table S3). Raw data are available in the GEO repository with the accession number GSE113272.

### Evolutionary conservation

Human orthologues of zebrafish genes (Table S4) were obtained using BioMart data mining tool from Ensembl. Next, the conservation scores from 100-way PhastCons^48^ conservation tracks were extracted using GENCODE (v.30) human exon coordinates corresponding to selected orthologues of zebrafish genes. PhastCons conservation 100 conservation tracks contain scores for the human genome calculated from multiple alignments with other 99 vertebrate species.

### KEGG and GO enrichment analysis

Gene Ontology (GO) and Kyoto Encyclopaedia of Genes and Genomes (KEGG) enrichment analysis for Illumina RNA sequencing data was performed using KEGG.db (v.3.2.3) and GO.db (v.3.4.1) R/Bioconductor packages. The KEGG enrichment analysis was done using zebrafish data from package org.Dr.eg.db (v 3.4.1). Pathways and genes detected in each fraction were filtered after Benjamini-Hochberg correction for an adjusted P-value < 0.05.

### Motif discovery

Homer^49^ and STREME^50^ tools were employed to extract motifs in the DNA sequence of 3’ and 5’UTR regions of genes enriched in MB and HS fractions in wild type and *chchd4a^-/-^* samples, respectively. Differential motif discovery using Homer software was performed using *findMotifsGenome.pl* script with default parameters, except –size 200 and –len 10. Whereas, analysis employing STREME was done using MEME Suite (v 5.4.1) and shuffled input sequences as a reference.

### RNAscope and immunostaining

Whole-mount fluorescent *in situ* hybridization by RNAscope was performed using RNAscope Multiplex Fluorescent v2 Assay kit (Advanced Cell Diagnostics) according to the manufacturer’s guideline with the following changes. 5dfp larvae in the *Tg(Xla*.*Eef1a1*:*mlsEGFP)cms1* were fixed with 4% paraformaldehyde for 18hrs at RT. The larvae were depigmented using a solution of 3% H_2_O_2_ and 1% KOH in dH_2_O for 10 min at RT. Further, the samples were washed in 1XPBS + 0.1% Tween20 (PBST), serially dehydrated and stored in 100% methanol at −20℃ until use. Next, the samples were treated with 0.2 M HCl in 100 % methanol for 10 min at RT and rehydrated in decreasing concentrations of methanol in 1xPBST. Upon final wash with PBST +1% BSA, the samples were transferred to a preheated target retrieval solution and heated in a thermoblock for 15 min at 98℃. Immediately the samples were immersed in PBST +1% BSA for 1 min, washed with 100% methanol and then again suspended in PBST +1% BSA. Further protease digestion was performed by treating the samples with Protease Plus reagent for 15 min in a preheated water bath at 40℃. This step was followed by the probe hybridization for 2h at 40℃. Hybridization with AMP1, AMP2 and AMP3 solutions and development of HRP signal were done as per the protocol. Samples were treated with a single probe at a time and in combination with TSA Plus Cyanine5 (PerkinElmer) at a dilution of 1:1500. We used *myod1* probe and custom-made *mt-nd5*, *rcn2*, *slc16a1a*, *comtd1* probes (Advanced Cell Diagnostics). Owing to the quenching of endogenous GFP signal, the RNAscope protocol was followed by immunostaining with anti-GFP antibody (GTX113617). Briefly, the samples were washed with PBST and blocked with a buffer containing 10% goat serum, 1% DMSO, 0.5% Triton X-100 in PBST for 1hr at RT with gentle shaking. The primary antibody was diluted 1:200 in the blocking buffer and the samples were incubated overnight at 4℃. Next, the samples were washed thrice with PBST for 10 min at RT and incubated with Alexa Fluor 488 coupled donkey anti-rabbit secondary antibody (ThermoFisher Scientific) diluted 1:500 in blocking buffer, for 1hr at RT. After washing thrice with PBST for 10 min at RT, the DNA was counterstained with DAPI (4′,6-diamidino-2-phenylindole, ThermoFisher Scientific) diluted 1:10,000 with PBST, for 40 min at RT. The larvae were mounted on glass slides with specially created cavities using Prolong Gold Antifade mountant (ThermoFisher Scientific). Images were acquired using LSM800 confocal laser scanning microscope (Zeiss). Region of interest defined by a dimension of 700 × 250 µm surrounding DAPI staining was chosen from three different biological replicates and Mander’s M1 colocalization coefficient defining the extent of mRNA fraction colocalized with mitochondria was calculated with the Manders Coefficients ImageJ plugin.

## Results

### Development and validation of subcellular fractionation method

To investigate which nuclear-encoded mitochondrial transcripts are likely to be located on the surface of mitochondria and hence participate in localized translation (Figure 1A), we established a new protocol for biochemical fractionation (Figure 1B) in zebrafish larvae. Although this approach largely adopts a strategy that was employed to obtain intact mitochondria with associated ribosomes in yeast^17^, it was optimized to obtain a membrane-bound fraction containing mitochondria and ER together with their associated ribosomes. Preservation of ER in this fraction allows us to perform more global analysis of membrane associated nuclear-encoded mitochondrial transcripts, as ER and mitochondria are highly interconnected in the cell. This not only includes their physical connections through mitochondria-associated membranes (MAMs)^51^, but also their extensive interactions supporting mitochondrial protein import^34–38^.

**Figure 1.**
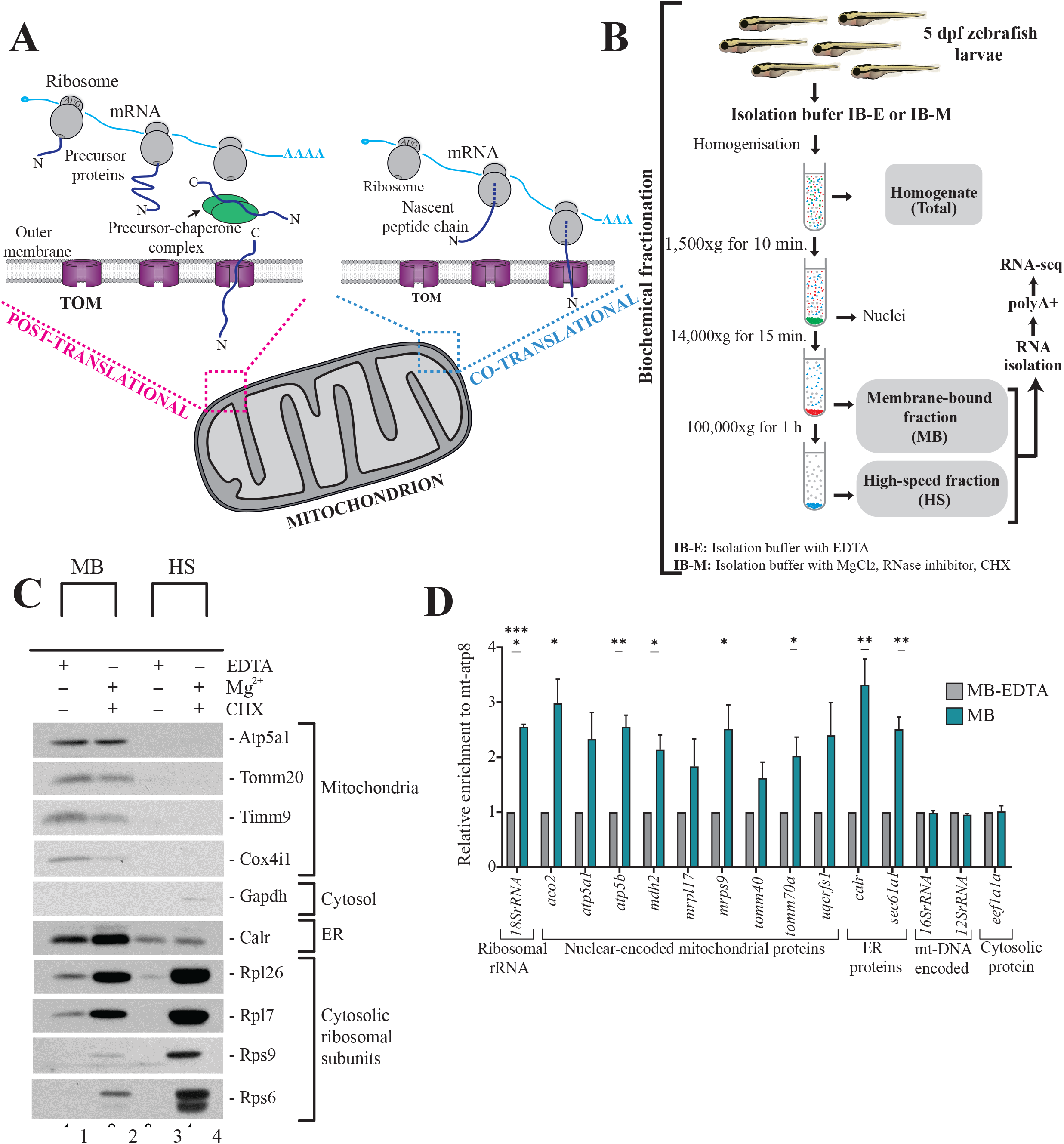
Isolation and identification of membrane associated mRNAs. (A) Routes for mitochondrial protein import. In post-translational import, proteins are imported to mitochondria after their complete synthesis in the cytosol in a process aided by chaperones and co-chaperones. Whereas, the localized translation that occurs at the surface of mitochondria may be directly coupled with the translocation of proteins into organelle. (B) Biochemical fractionation of 5dpf zebrafish larvae based on strategy to obtain intact mitochondria with associated ribosomes in yeast^17^. Isolation of mitochondria with associated ribosomes was performed using a ribosome-friendly buffer (IB-M) containing MgCl_2_, cycloheximide (CHX) and RNAse inhibitors. MB and HS fractions were obtained via differential centrifugation. Total RNA isolated from each fraction was enriched for polyadenylated fraction and subjected to short-read sequencing using Illumina platform. (C) Isolation of membrane-bound and high speed fractions. (D) RT-qPCR validation of MB fraction. Relative enrichment of genes in MB with respect to EDTA-stripped MB upon normalization with mitochondrial DNA encoded mt-atp8 is shown. Data derived from three biological replicates. Error bars correspond to SEM; *P <* 0.05 (*), *P <* 0.01 (**) and *P <* 0.0001 (****) by unpaired *t*-test.

Standard biochemical strategies for isolating mitochondria utilize EDTA as one of the protease inhibitors in the isolation buffer (IB-E). EDTA disrupts association of cytosolic ribosomes with mitochondria by chelating the metal ions that keep the ribosomes intact. Therefore, for the isolation of mitochondria with associated ribosomes, a ribosome-friendly buffer (IB-M) containing MgCl_2_, cycloheximide (CHX) and RNAse inhibitors was used. Magnesium ions are essential for the integrity of ribosomes^52^, while CHX stabilizes ribosome-nascent chain complexes. By homogenization and differential centrifugation using IB-E or IB-M, we obtained three subcellular fractions – membrane-bound (MB), high-speed (HS) and whole, unfractionated cell (total) (Figure 1B). Analysis of subcellular fractionation by western blotting showed that the membrane-bound fraction was exclusively enriched with intact mitochondria as indicated by the mitochondrial markers, including proteins from various subcompartments - Tomm20 of outer mitochondrial membrane, Cox4i1 of inner mitochondrial membrane and Timm9 of intermembrane space (Figure S1E, lane 1-3). These markers were not visible in the high-speed and whole-cell fractions. The ER marker Calreticulin (Calr) was also enriched in the membrane-bound fraction compared to unfractionated cells and high-speed fraction obtained using both buffer systems (Figure S1E, lane 4). Interestingly, parts of ER were also detected in the HS fraction, which means that due to its size and diverse structure, ER splits between two fractions. However, the HS fraction showed no enrichment of mitochondrial markers. Additionally, both high-speed and membrane-bound fractions obtained with IB-M were enriched in ribosomal proteins (Figure S1E, lane 5 and 6). In contrast, the cytosolic fraction obtained with IB-M did not show any ribosomal protein enrichment. We reasoned that this could be due to the co-sedimentation of free polysomes with HS fraction at very high speed used during ultracentrifugation. Consequently, the HS fraction contains a combination of both ER-bound ribosomes and free cytosolic polysomes. We also noticed that Gapdh, the cytosolic protein, was mainly enriched in the cytosolic fraction (Figure 1C, lane 7).

After determining the composition of subcellular fractions, we isolated MB and HS fractions with IB-M containing BSA. Addition of BSA in IB-M prevents any proteolysis of mitochondria that might occur during fractionation and enables reliable profiling of MB-associated RNAs. The comparison between the IB-M isolated and the EDTA-stripped MB fractions (isolated with IB-E containing BSA), once again confirmed the presence of intact mitochondria and enrichment of cytosolic ribosomal subunits in the IB-M isolated fraction (Figure 1C). Analysis of the HS fraction also confirmed the enrichment of cytosolic ribosomes in comparison to its EDTA-stripped counterpart. Additionally, an abundant presence of ER in the IB-M isolated MB was also noticed in comparison to the EDTA-stripped MB fraction. Whereas, the IB-M isolated and EDTA-stripped HS fractions showed similar ER enrichment. The IB-M isolated MB and HS fractions were used for all subsequent experiments, and henceforth we will refer to them as MB and HS.

To prove that the MB fraction is enriched with nuclear-encoded mitochondrial transcripts, relative RNA levels of the MB with respect to the EDTA-stripped MB were determined by the RT-qPCR analysis (Figure 1E). This analysis included mainly zebrafish orthologues of nuclear-encoded mitochondrial transcripts known to be localized on the surface of mitochondria in yeast^20^. Five out of nine of these nuclear-encoded mitochondrial mRNAs were significantly enriched in the MB with respect to its EDTA-stripped counterpart. Whereas, the levels of mt-DNA encoded transcripts and the *eef1a1a* transcript encoding a cytosolic protein remained similar between MB and EDTA-stripped MB fraction. In line with the previous observations pointing towards the enrichment of ER fragments in the MB, the *sec61a1* and *calr* transcripts encoding ER membrane and the ER luminal proteins, respectively, were also found to be enriched in the MB fraction. We also confirmed the enrichment of 18S rRNA – the structural component of the cytosolic ribosome in this fraction.

### High-throughput profiling of subcellular localization of mitochondrial gene products

To examine the subcellular localization of nuclear-encoded mitochondrial transcripts, we isolated polyadenylated RNA from MB and HS fractions obtained via biochemical fractionation from wild type zebrafish individuals. We used short-read Illumina RNA sequencing to identify genes with transcripts enriched in each fraction. Interestingly, we observed two populations of genes (Figure S2A) enriched either in HS or MB. We did not detect genes that would be present to the same extent in both fractions. One of the possible reasons for this, could be exclusion of poorly enriched genes from the analysis (see Materials and Methods for details). In total, the differential analysis showed enrichment of 2,177 and 3,315 genes in MB and HS fractions, respectively (Figure 2A). To reliably assign genes to each fraction, we required at least 2-fold enrichment (highly enriched) between MB and HS fractions (FDR 5%; log2FC <=> +/− 1). As this threshold was selected arbitrarily to match the common standards of RNA-seq data analysis, we also distinguished a class of moderately enriched genes (FDR 5%; 0.5 =< log2FC < 1 or −1 < log2FC <= −0.5). However, this group was much less numerous with 1,455 and 1,316 genes enriched in MB and HS fractions (Figure 2A). We also observed a substantial enrichment of all 13 genes encoded by the mitochondrial genome among the MB highly enriched genes (Figure S2B), which confirms the presence of mitochondria in this fraction.

**Figure 2.**
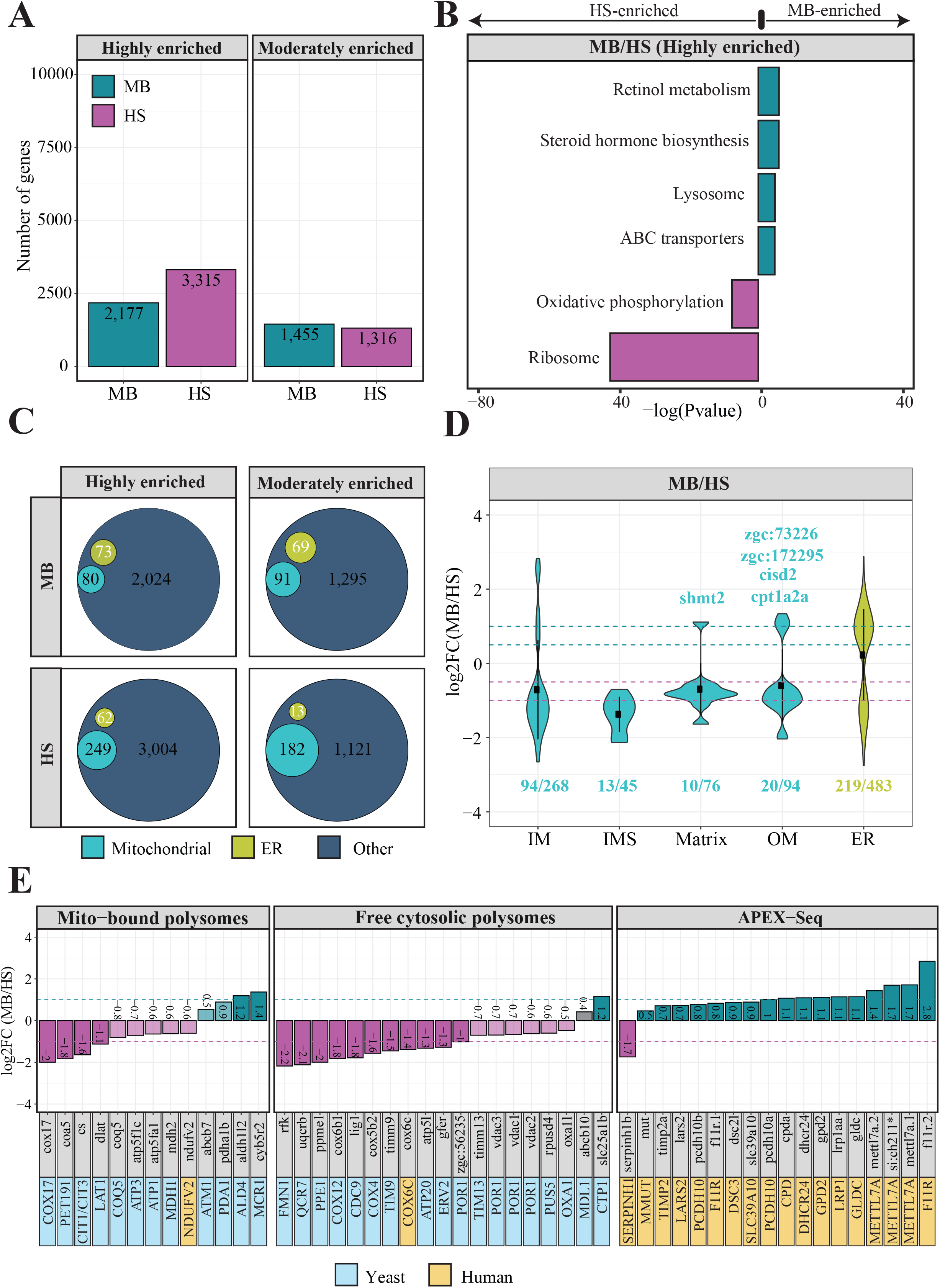
Asymmetric subcellular localization of nuclear-encoded mitochondrial mRNAs. (A) Bar plots representing the total number of genes identified in MB and HS fractions obtained via biochemical fractionation of wild type 5 dpf zebrafish samples and short-read Illumina RNA sequencing. (B) Kyoto Encyclopaedia of Genes and Genomes (KEGG) enrichment analysis using highly enriched 2,177 and 3,315 genes. The results are shown as a negative log10 P-value after Bonferroni correction. Bars in green indicate KEGG terms enriched for genes detected in MB fraction, while the purple bars represent the KEGG terms for the features identified in HS. For better representation of the results the bars enriched for genes in HS were transformed to be located on the left hand side of the plot. (C) Venn diagrams representing the number of mitochondrial genes from the merged MitoCarta 2.0^55^ and IMPI^56^ repository, as well as the zebrafish orthologues of human 483 genes encoding proteins that localize to ER according to the Human Protein Atlas, identified in MB or HS fraction from WT samples. (D) Violin plot showing the distribution of log_2_ gene enrichments for genes grouped by their location within mitochondria. ER genes were also included in this analysis (E) Bar plots showing the log_2_ gene enrichments of yeast (blue) and human (yellow) orthologous genes for which transcripts were reported to be translated by the mitochondrion-bound or free cytosolic polysomes.

To determine the function of genes assigned to each fraction, we performed an enrichment analysis using the Kyoto Encyclopaedia of Genes and Genomes (KEGG). This analysis revealed highly enriched genes in the HS fraction to be either involved in the oxidative phosphorylation (OXPHOS) or encoding ribosomal proteins (Figure 2B). Interestingly, KEGG analysis also reported a link between the MB fraction and ATP-binding cassette (ABC) transporters. No enriched KEGG pathways were detected for moderately enriched genes in MB and HS fractions.

Next, we carefully investigated the subcellular localization of mitochondrial gene products by using merged set of zebrafish orthologues for two independent human gene inventories with strong support of mitochondrial localization: MitoCarta 2.0^53^ and Integrated Mitochondrial Protein Index (IMPI)^54^. This analysis revealed 80 and 249 mitochondrial genes to be highly enriched in the MB and HS fractions, respectively (Figure 2C and S2C), indicating that nuclear-encoded mitochondrial genes are mainly present in the HS fraction. We noticed the same trend for moderately enriched genes with 91 and 182 mitochondrial genes detected in MB and HS fractions, respectively (Figure 2C and S2D). To investigate the enrichment of ER genes in each fraction, we used zebrafish orthologues of 483 (∼2% of all protein coding human genes) human genes that encode proteins localizing to the ER (Table S5), as shown by the Human Protein Atlas^55^. Our results indicate that the ER genes are uniformly enriched across all fractions, with an exception of only 13 detected in the moderately enriched gene set for the HS fraction.

To further explore the presence of mitochondrial genes in HS and MB fractions, we analyzed the distributions of enrichment values for different mitochondrial compartments (Figure 2D). Most genes for each mitochondrial compartment have their transcripts enriched in the HS, whereas individual cases like *shmt2* gene (matrix) involved in the one-carbon pathway^56^ are enriched in the MB fraction. The transcripts encoding IMS proteins are distinct from others subgroups, as they were almost entirely localized in the HS fraction. Importantly, the MB enrichment observed for the IM proteins is solely driven by the enrichment of mitochondrially encoded genes (Figure S2B). According to our previous observations, ER genes appeared to be uniformly enriched across investigated fractions, which is in line with estimated fraction composition (Figure 1C).

We further compared observed gene enrichments with the outcome of other studies investigating the subcellular localization of mitochondrial gene products, mainly in yeast^3,9,14^ and human cell lines^28^. Using our data, we analyzed the enrichment of genes with transcripts known to be located in the proximity of mitochondria, translated by mitochondria-bound or free cytosolic polysomes. Prior to this analysis, we identified a zebrafish orthologue for each of those genes. Interestingly, only two yeast genes *ALD4* and *MCR1* for which transcripts are known to be translated by the mitochondria-bound polysomes^9,14^ were highly enriched, while the other two *ATM1*^12^ and *PDA1*^14^ were moderately enriched in the MB fraction (Figure 2E). On the other hand we observed a much higher validation rate for genes known to be translated by the free cytosolic polysomes, as 11 and 6 out of 19 investigated genes showed high and moderate enrichment in HS fraction, respectively. To clarify the poor overlap between our membrane-associated transcripts and those translated by the mitochondria-bound polysomes in yeast, we explored the enrichment of zebrafish orthologues of human genes, which had their products detected in the close proximity of OM by APEX-Seq study^28^. In this comparison, 94% (17/18) genes were also enriched in our MB fraction (Figure 2E, panel 3). This could suggest that the discrepancy between the enrichment of transcripts located at the surface of mitochondria in zebrafish and yeast is driven by the differences in the organismal complexity. In particular, we observed poor correlation between the MB enrichment values for zebrafish orthologues in our study and the mitochondrial enrichment reported by the high-throughput proximity specific ribosome profiling in yeast with and without CHX treatment^20^ (Figure S3).

### Properties of membrane-associated mitochondrial transcripts

To understand the principles determining the specific cellular localization of transcripts encoding mitochondrial proteins, we investigated the enrichment of specific gene families. We started this analysis by looking at ABC transporters, as suggested by the KEGG pathway enrichment analysis (Figure 2B). The ABC transporters are ubiquitous membrane-bound proteins that are also ATP-dependent pumps, allowing to move substrates in or out of cells^57^. Although ABC transporters can be divided into seven different subfamilies (designated A to G), for our analysis, we distinguished only mitochondrial and non-mitochondrial subgroups of ABC transporters. All genes coding for mitochondrial ABC transporters (B subfamily) were moderately enriched in the MB fraction, with the *ABCB10* showing the smallest enrichment (log_2_FC = 0.42) (Figure 3A). The genes encoding non-mitochondrial ABC transporters showed high enrichment in the MB fraction, with the *ABCG5* being the most highly enriched (log_2_FC = 2.82) gene. We also observed genes encoding non-mitochondrial solute carriers (SLC) to be more abundant in the MB fraction over HS, whereas genes coding for mitochondrial solute carrier family (SLC25) were enriched in HS fraction (Figure 3B). The solute carriers are membrane-bound proteins regulating transport of various substances, including organic cations, ions and amino acids over the cell membrane. The mitochondrial solute carriers transport nutrients required for the energy conversion and maintenance of the cell across the mitochondrial inner membrane^58^. One of the most notable common features among ABC transporters and solute carriers is the presence of hydrophobic transmembrane domains (TMDs). To fulfil their function, proteins with TMDs must protect themselves from random integration into other membranes and aggregation in the cytosol. Therefore, the localized translation coupled with membrane insertion could help to reduce the problem of mis-targeted membrane integration and accumulation of proteins with TMDs in the cytosol. ER can act as the platform for the co-translational import and segregation of proteins with TMDs^59,60^. Thus, the enrichment of non-mitochondrial ABC transporters and solute carriers in the MB fraction is largely ER driven.

**Figure 3.**
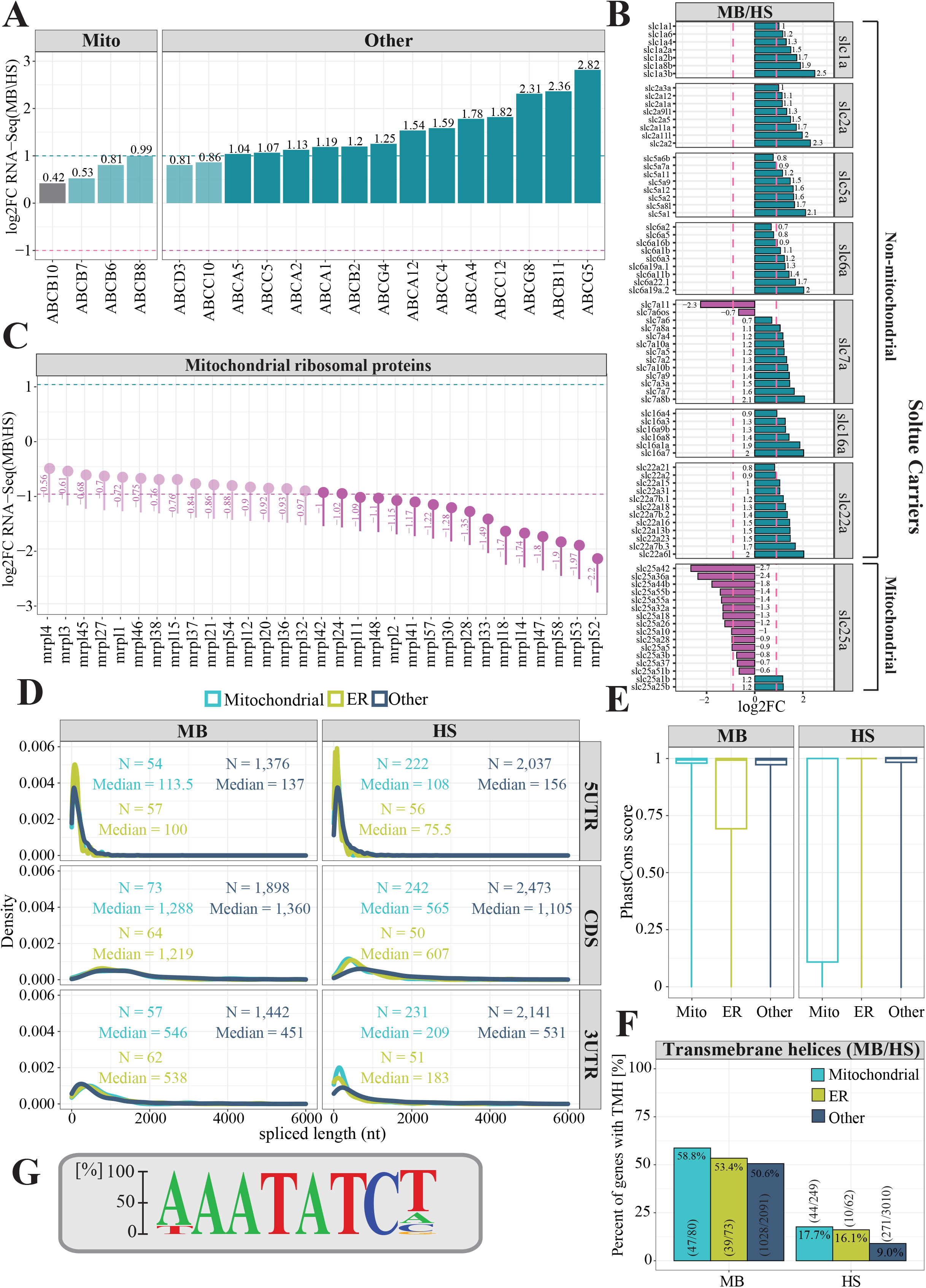
Properties of membrane-associated transcripts. Analysis of gene enrichment of (A) the ATP-binding cassette transporters (ABC transporters), (B) Solute carriers and (C) Mitochondrial ribosomal proteins. The purple dashed line represents log_2_FC = −1 and the green dashed line represents log_2_FC = 1. (D) Length distribution analysis of coding sequences (CDS) and UTR regions for transcripts encoding mitochondrial (cyan), ER (green) and other proteins (navy). (E) Boxplots showing PhastCons^48^ conservation tracks for gene categories described above. PhastCons score equal to 1, represents highly conserved sequences, while PhastCons score equal to 0 indicates rapidly evolving sequences. (F) Bar plot representing proportion of transcripts encoding mitochondrial, ER and other proteins with transmembrane domains. The numbers in parenthesis indicate figures used to calculate proportions (G) The sequence logo illustrating motif enriched in 3’UTR regions of MB fraction-enriched transcripts encoding mitochondrial proteins.

We also noticed transcripts encoding mitochondrial ribosomal proteins to be entirely enriched in the HS fraction (Figure 3C). This is consistent with previous findings, showing that mitochondrial ribosomal proteins as soluble proteins are mainly synthetized in the cytosol and imported into mitochondrial matrix in their precursor form^9^.

We further explored the properties of transcripts enriched in each fraction by comparing their lengths, particularly of their 5’ and 3’ untranslated regions (UTR) and coding sequences (CDSs). Surprisingly, CDSs of transcripts encoding mitochondrial proteins that were enriched in MB are two times longer (median = 1,288 nt) compared to the ones enriched in the HS fraction (median = 565 nt, P=2.51e-11, Wilcoxon rank sum test with continuity correction). This is also true to the same extent for ER transcripts (1,219 nt vs. 607 nt, P=3.23e-12) and to a lower extent for other transcripts, as MB enriched CDSs are ∼30% longer than those for transcripts detected in HS fraction (P=2.2e-16, Wilcoxon rank sum test with continuity correction) (Figure 3D). Moreover, transcripts encoding mitochondrial and ER proteins that have been identified in the MB fraction had on average much longer 3’UTRs (median = 546 nt and 538 nt) with respect to the HS fraction enriched transcripts (median = 209 nt and 183 nt, P=5.24e-06 and P=2.32e-09, respectively Wilcoxon rank sum test with continuity correction). Whereas, the length of 3’UTRs of other transcripts did not vary that much across fractions (P=8.76e-05, Wilcoxon rank sum test with continuity correction). Next, we investigated the evolutionary conservation of mitochondrial, ER and other transcripts enriched in the MB and HS fractions using a 100-way vertebrate sequence alignment^48,61^. This analysis revealed that the enriched transcripts in the MB fraction, with exception to the ER ones, were highly conserved, while the ones present in the HS fraction were more rapidly evolving (Figure 3E). This is particularly true for transcripts encoding mitochondrial proteins and is consistent with previous findings, indicating that transcripts translated by mitochondrion-bound polysomes are in general much more evolutionary conserved^14,62,63^. Our results also confirmed that all types (mitochondrial, ER and other) of membrane associated transcripts were on average likely to encode proteins with transmembrane domains (58.8% vs. 17.7%, 53.4% vs. 16.1% and 50.6% vs. 9%) (Figure 3F). Finally, the analysis of 3’UTR regions of mitochondrial transcripts detected in the MB fraction using Homer^49^ software revealed the presence of 16 enriched sequence motifs (Figure S4A). One motif among them – AAATATCY, where Y in 60% of cases is substituted by T (Figure 3G) was clearly enriched. This motif was also detected in 3’UTR regions of MB enriched mitochondrial transcripts by the STREME – a recently developed software for motif discovery^50^. Interestingly, among known motifs similar to this one, STREME indicated the Puf3 protein binding motif (TGTAAATA)^64^. Puf3 associates with the cytosolic surface of the OM and guides mRNAs to the mitochondrial surface through the recognition of the binding motif in yeast^65^. The motif comparison performed by Tomtom software^66^ revealed overlap of 6 nucleotides between motif sequences (Figure S4B). The FOXJ3 – a human transcription factor was another motif detected as similar by the STREME software. These two motifs share 11 nucleotides (Figure S4C). According to STREME, presented motif was detected in 32.2% (p-value = 7.53e-001, E-value=2.3e+000) of analyzed transcript sequences.

### Experimental validation of predicted subcellular locations

To validate the computationally predicted enrichment of mitochondrial mRNAs, RT-qPCR experiments using MB and HS fractions were performed. For the validation of MB enriched mitochondrial transcripts (Figure 4A), three candidates from highly enriched category and three from moderately enriched category were selected (Figure 4B). For all selected genes (*comtd1*, *slc16a1a*, *cyb5r2*, *abcb6a*, *rcn2* and *chpt1*), we were able to confirm their enrichment in the membrane-bound fraction. As expected, *mt-atp8* encoded by the mitochondrial genome has also shown substantial MB enrichment. For the validation of genes enriched in the HS fraction (Figure 4C), we selected candidates encoding different classes of mitochondrial proteins such as Mia40 substrates, tail-anchored (TA) and mitoribosomal proteins. The RT-qPCR results showed that mRNAs encoding mitochondrial ribosomal proteins, Mia40 substrates and TA mitochondrial proteins are enriched in the HS fraction (Figure 4D). Therefore, in all cases, the RT-qPCR results were in the agreement with the RNA-seq data (Figure 4A and C).

**Figure 4.**
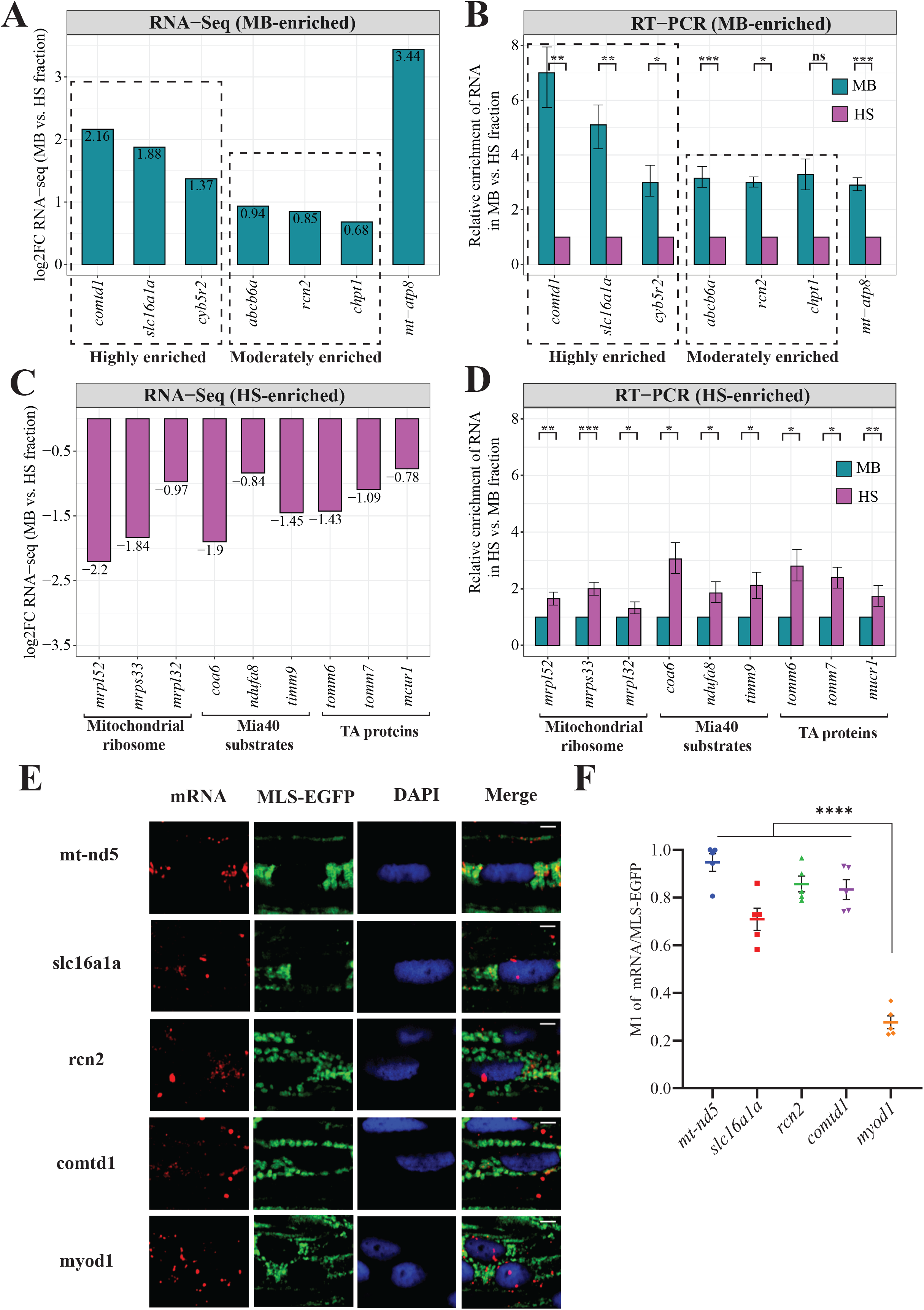
Experimental validation of computationally predicted gene enrichments. (A) Bar plot showing the log_2_ gene enrichments for the MB fraction-enriched MitoCarta 2.0 genes detected as highly and moderately enriched. (B) RT-qPCR validation of MB fraction-enriched MitoCarta 2.0 genes. (C) Log_2_ gene enrichments for the HS fraction-enriched MitoCarta 2.0 genes encoding mitochondrial ribosomal proteins, Mia40 substrates and TA proteins. (C) RT-qPCR validation of genes enriched in the HS fraction. Relative enrichment of genes in MB with respect to HS upon normalization with ERCC-00096 is shown. Data derived from three biological replicates. Error bars correspond to SEM; *P <* 0.05 (*), *P <* 0.01 (**), *P <* 0.001 (***) and *P <* 0.0001 (****) by unpaired *t*-test (C) Fluorescent *in situ* hybridization of selected candidates by RNAscope on whole mount 5dpf zebrafish larvae. mRNAs are probed by specific RNAscope probes (red) and mitochondria are stained with anti-GFP antibody (green). Yellow in the merged image indicates co-localization of mRNAs on MLS-EGFP tagged mitochondria. Scale = 5µm (D) Analysis of co-localization of MB fraction-enriched MitoCarta2.0 mRNAs with mitochondria. M1 indicates Mander’s co-localization co-efficient of mRNA fraction co-localized on mitochondria harboring MLS-EGFP. The difference between *myod1* negative control and mRNA candidates were analyzed by unpaired *t*-test. Error bars correspond to SEM and data derived from five region of interests originating from three experiments (n=5); *P <* 0.0001 (****).

To prove that MB enriched mitochondrial transcripts are actually present on the surface of mitochondria *in vivo*, we performed fluorescent *in-situ* hybridization by RNAscope technique^67,68^. We selected three MB fraction enriched genes - *rcn2*, *slc16a1a*, and *comtd1,* which are zebrafish orthologues of human genes present in MitoCarta 2.0 gene set^53^. The transcripts of *rcn2* and *slc16a1a* have been recently reported by high-throughput APEX-seq study to be associated with outer mitochondrial membrane in human cells^28^. Proteins encoded by these three genes were also showed to be localizing to mitochondria by numerous studies^69–74^. As expected, the mtDNA encoded mRNA, *mt-nd5*, presented the highest co-localization with mitochondria (Figure 4E and F). Transcripts of three genes *rcn2*, *slc16a1a*, and *comtd1* enriched in the MB fraction also displayed high mitochondrial localization. In contrast, *myod1* mRNA, which was not enriched in the MB fraction according to our analysis, presented a cytosolic distribution and the lowest co-localization with mitochondria. Therefore, our RNAscope results indicate that the transcripts of nuclear-encoded mitochondrial genes enriched in MB fraction, show high degree of localization on the mitochondrial surface in zebrafish.

### Transcriptome changes triggered by the *chchd4a^-/-^* mutation

Deficiency in any mitochondrial import machinery results in the accumulation of mitochondrial precursor proteins in the cytosol that triggers the defensive response^31^. Cells protect themselves against defects in mitochondrial biogenesis by both inhibiting protein synthesis and activating the proteasome that performs cellular protein clearance. Since proteasomal activity is regulated by the quantity of mis-localized mitochondria-destined precursor proteins, we hypothesize that defects in the mitochondrial import pathway could alter the localization of mRNAs and translation of nuclear-encoded mitochondrial proteins at the surface of mitochondria to reduce the number of over-accumulated precursor proteins in the cytosol.

To explore this hypothesis, we took advantage of *chchd4a*^-/-^ zebrafish mutant line, in which the import of mitochondrial proteins into IMS is disrupted^39^. Mutation in *chchd4a* triggers a glycolytic phenotype in zebrafish, leading to starvation and death starting at 10dpf. To examine in details the stress response caused by *chchd4a^-/-^* mutation, we re-analyzed the RNA sequencing data from our previous study, where we directly compared *chchd4a^-/-^* to the wild type 5dpf zebrafish samples^39^. In total, the differential analysis showed expression changes for 217 genes (FDR 5%; log2FC +/− 0.9) with 110 and 107 being up- and down-regulated, respectively (Figure 5A). In general, we observed increased expression of genes involved in amino acid activation and protein refolding (*hsp70, hspd1, hspa9, hspa5*) (Figure 5B). Interestingly, we also noticed four genes from the one-carbon pathway (*shmt2, mthfd2, aldh1l2, mthfd1l)*, whose proteins are known to localize to mitochondria, to be upregulated by *chchd4a^-/-^* mutation (Figure 5C). We previously observed elevated expression of these genes also at the protein level upon *chchd4a^-/-^* mutation^39^. It has been shown that the one-carbon metabolism can produce mitochondrial NADH under conditions of stress, when the TCA cycle affected by the cell conditions loses its capacity^56^. Together with the increased expression of genes encoding mitochondrial and cytoplasmic one-carbon pathway enzymes, we also observed transcriptional activation of serine metabolism, in particular L-serine biosynthetic process (*phgdh* and *psat1*). This is in line with previous observations showing that serine is driving the NADH production via one-carbon metabolism and that various stress responses can increase the transcription of enzymes involved in serine biosynthesis^75,76^. Increased transcription of one-carbon metabolism genes suggests a change in the way NADH is produced in the cell.

**Figure 5.**
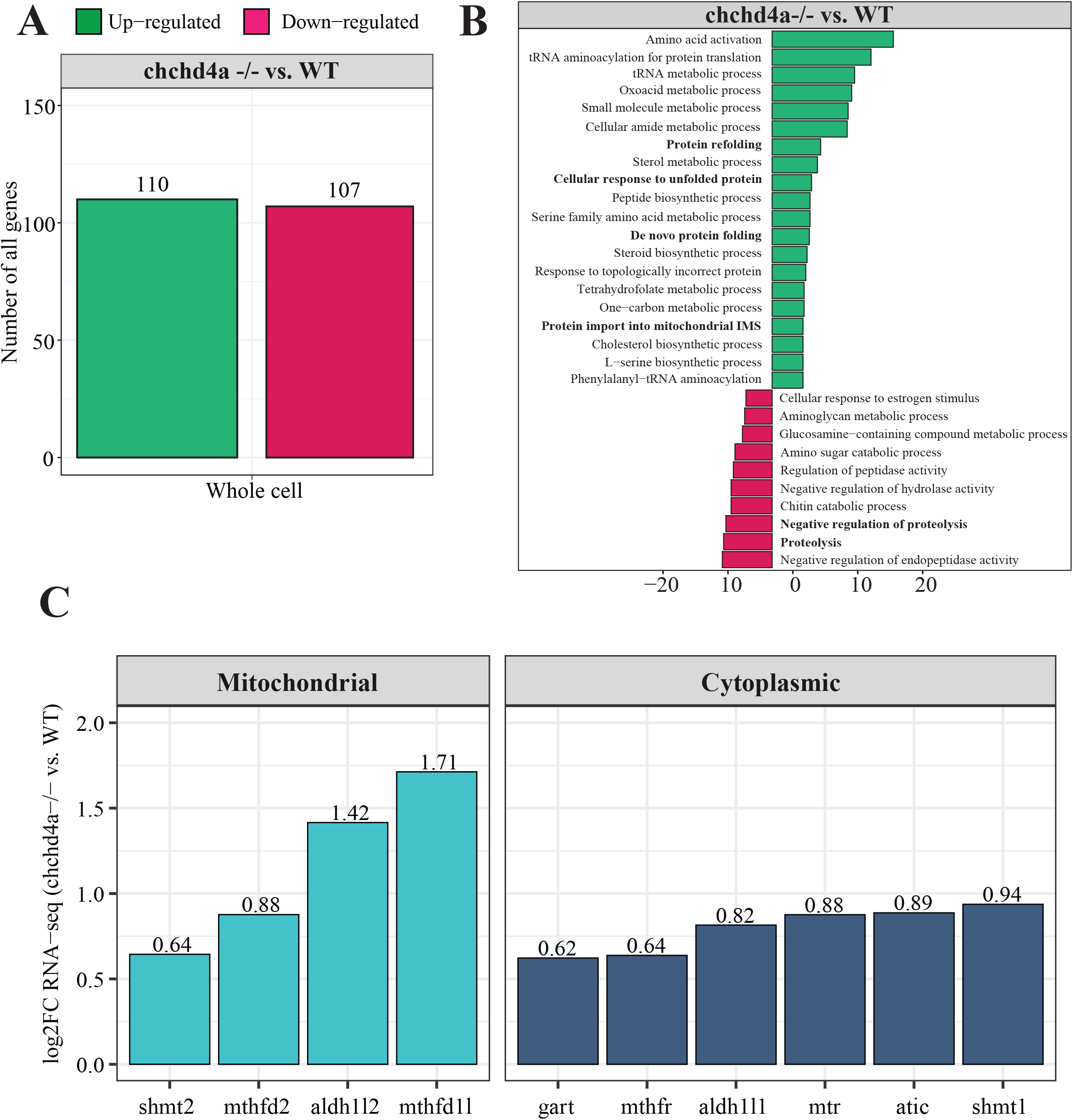
Transcriptome changes triggered by *chchd4a^-^/^-^* mutation. (A) Numbers of differentially expressed genes between whole cell unfractionated *chchd4a^-^/^-^* and wild type samples. (B) KEGG enrichment analysis for differentially expressed genes for the whole unfractionated samples. (C) Bar plot showing expression changes of one-carbon genes encoding cytoplasmic (dark blue) and mitochondrial enzymes (light blue).

### Dynamics of transcript subcellular localization under mitochondrial stress conditions

To explore the impact of *chchd4a*^-/-^ mutation on the subcellular localization of nuclear-encoded mitochondrial transcripts, we obtained MB and HS fractions via biochemical fractionation from 5dpf *chchd4a*^-/-^ zebrafish larvae (Figure S5). Next, we isolated polyadenylated RNA from each fraction and performed RNA-seq. We followed the same data analysis steps, in which we directly compared the MB to the HS and assigned genes to each fraction based on their enrichment (FDR 5%; log2FC <=> +/− 1). Again, we noticed two populations of genes either enriched in the MB or HS fraction (Figure S6A). No major changes in the number of detected genes in *chchd4a*^-/-^ compared to wild type (WT) samples were observed, except for the MB fraction, in which the number of enriched genes increased by almost a thousand (Figure S6B).

We also noticed that the *chchd4a*^-/-^ mutation affected the enrichment of mitochondrially encoded genes in the MB fraction, particularly for three genes: *mt−nd3*, *mt−nd6* and *mt−atp6* (Figure S6C). To determine the function of genes identified in each fraction we performed a KEGG enrichment analysis. We noticed further enrichment of genes involved in the oxidative phosphorylation, together with genes encoding cytosolic and mitochondrial ribosomal proteins in the HS fraction compared to wild type (Figure S6D). In contrast, genes detected in the MB fraction were mainly enriched in general KEGG terms, including lysosome, sphingolipid metabolism or ECM−receptor interaction. Similar to the analysis strategy for the wild type, we again tested our gene lists against a merged set of zebrafish orthologues for genes in MitoCarta 2.0^53^ and IMPI^54^ inventories. Indeed, we observed an increased presence of nuclear-encoded mitochondrial genes in the HS fraction for both highly and moderately enriched categories, while in the MB there is a slight increase in the number of mitochondrial transcripts in the highly enriched category of *chchd4a*-/- mutants as compared to wild type (Figure 6A). This was also clearly seen in the distribution of enrichment values for specific mitochondrial compartments (Figure 6B) by a noticeable shift towards HS fraction for genes from all the mitochondrial sub-localizations. To understand if this shift was directly caused by the *chchd4a*^-/-^ mutation, we investigated the enrichment of MIA pathway targets. This analysis confirmed that the *chchd4a*^-/-^ mutation, indeed, enhanced the HS fraction enrichment for almost all of analyzed MIA pathway targets (Figure 6C).

**Figure 6.**
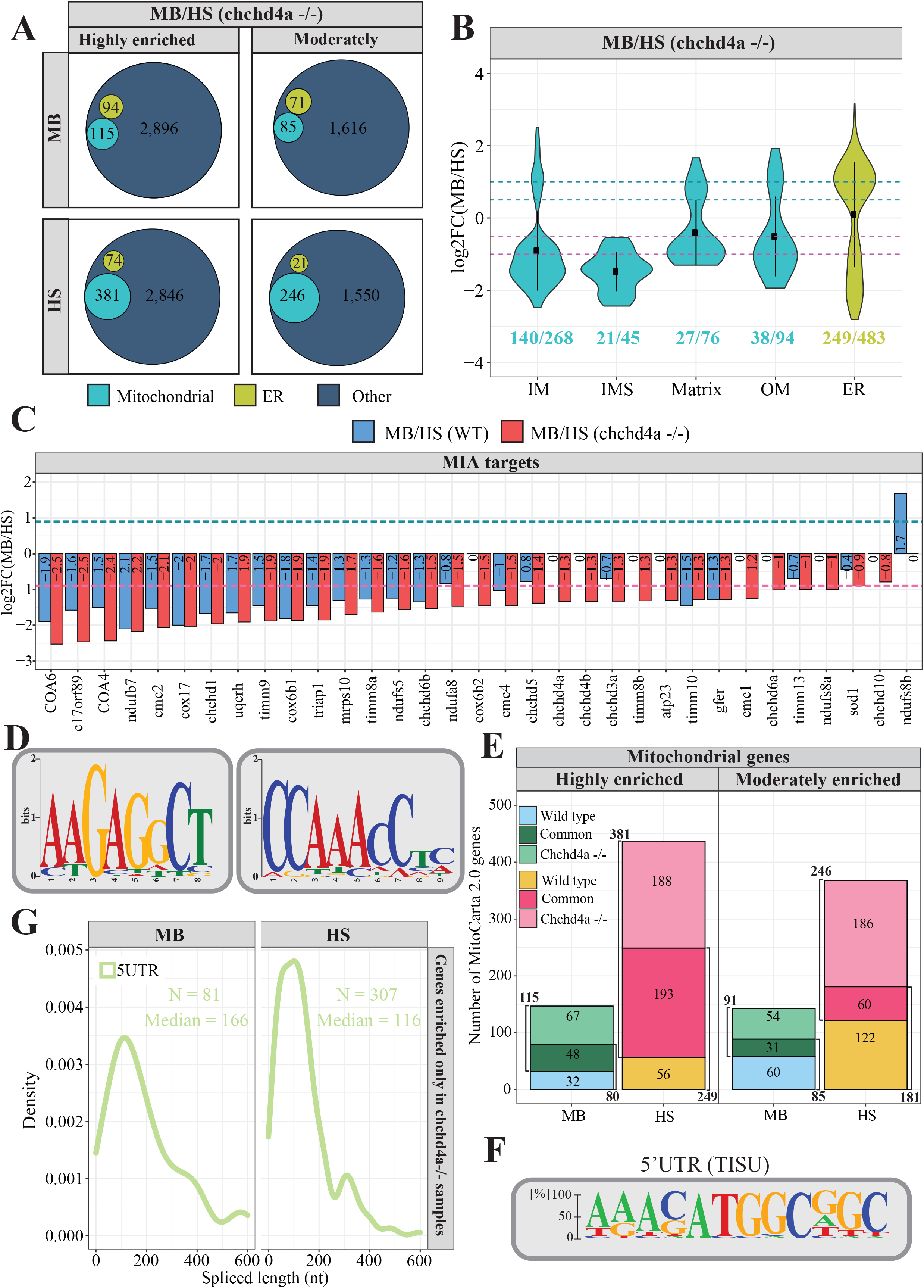
Dynamics of mRNA subcellular localization under mitochondrial protein import deficiency. (A) Venn diagrams representing the number of mitochondrial genes from the merged MitoCarta 2.0^55^ and IMPI^56^ repository, as well as the zebrafish orthologues of human 483 genes encoding proteins that localize to ER according to the Human Protein Atlas in each *chchd4a^-^/^-^* set. (B) Violin plot showing the distribution of log_2_ *chchd4a^-^/^-^* gene enrichments for genes grouped by their location within mitochondria, including also ER genes; (C) Gene enrichment analysis for MIA substrates in transcriptomic data for wild type (blue) and *chchd4a^-^/^-^* (red) 5dpf samples. The purple dashed line represents log_2_FC = −1 and the green dashed line represents log_2_FC = 1. (D) Two sequence logo illustrating motifs detected in 3’UTR regions of MB fraction-enriched transcripts encoding mitochondrial proteins in *chchd4a^-^/^-^* samples identified by STREME software^66^ (E) Venn diagrams depicting the overlap between the genes identified in wild type and *chchd4a^-^/^-^* MB and HS fractions, respectively. (F) The sequence logo illustrating TISU element found in the 5’UTR regions of transcripts exclusively enriched in the HS upon *chchd4a* mutation. (G) Distribution of 5’UTR lengths in which TISU element was detected.

To further investigate if mitochondrial protein import disorders had affected the properties of membrane-associated transcripts, we compared the transcript lengths between the MB and HS fractions in *chchd4a*^-/-^. Interestingly, mitochondrial, ER and other transcripts enriched in the MB fraction have on average even longer CDSs (Figure S6E), when compared to the wild type MB fraction (Figure 3D). The median length increased by ∼200 for mitochondrial and ER transcripts (P=2.2e-16 and P=3.67e-11 according to Wilcoxon rank sum test with continuity correction), while for ∼300 nucleotides for other types of transcripts (P=2.2e-15, Wilcoxon rank sum test with continuity correction). Moreover, we noticed that the mitochondrial transcripts in the MB fraction of *chchd4a*^-/-^ mutants had shorter 3’UTR regions (median = 424.5 nt) than those identified in wild type MB fraction (median = 546, P=0.00017 according to Wilcoxon rank sum test with continuity correction). This effect is less pronounced for the ER transcripts (median = 538 nt vs. median = 496 nt) Surprisingly, the length of CDSs in other types of transcripts enriched in the HS fraction was reduced by ∼40% (1,105 vs. 667, P=0.00023). This means that the average CDS length of transcripts enriched in the MB fraction are almost 3 times longer than those detected in the HS fraction. Thus, the difference between the CDS length for transcripts in MB and HS fractions and hence, the size of the protein, was even larger for *chchd4a*^-/-^ mutants compared to the wild type. Surprisingly, similar to that in wild-type, transcripts enriched in the MB fraction of *chchd4a*^-/-^ mutant were also more evolutionarily conserved than those in the HS fraction according to the analysis performed using 100-way Multiz vertebrate alignment^48,61^ (Figure S6F). Interestingly, we observed the biggest change in the evolutionary conservation scores for the mitochondrial transcripts. At the same time, *chchd4a*^-/-^ mutation reduced the presence of transcripts encoding proteins with TMDs in the MB fraction for both mitochondrial and other transcripts, whereas this proportion remained similar for ER transcripts (Figure S6G). Despite the reduced length of 3’UTRs of mitochondrial transcripts enriched in the MB fraction upon *chchd4a*^-/-^ mutation, using Homer^49^ software we confirmed the presence of previously detected sequence motif (AAATATCY, 26.69%, p-value=1e-2) (Figure 3G) among 15 detected motifs (Figure S7A). However, we were not able to detect this motif using STREME^50^ software (Figure S7B). Instead, STREME^50^ reported two new motifs enriched in 3’UTR regions of mitochondrial transcripts detected in the MB *chchd4a*^-/-^ fraction – AAGAGCCT and CCAAACCTC (Figure 6D). These motifs were found in 26.6% (p-value=5.0e-0001 and E-value=1.5e+000) and 35.4% (p-value=5.0e-0001 and E-value=1.5e+000) 3’UTR sequences of mitochondrial transcripts respectively. Although the mechanisms driving the localization of mitochondrial transcripts to the MB fraction upon *chchd4a*^-/-^ mutation remain unknown, our data suggests that the *chchd4a* loss of function further limits the type of localized mRNAs to well-conserved transcripts encoding large proteins.

To explore the increased enrichment of mitochondrial transcripts in the HS fraction, we further investigated the overlap between genes enriched in each fraction for wild type and *chchd4a^-/-^*. Our results indicate that 60% (48) and ∼80% (193) of highly enriched genes are shared between *chchd4a*^-/-^ and wild type MB and HS fractions, respectively (Figure 6E). This could suggest that the *chchd4a*^-/-^ mutation is further enriching the RNA subpopulations in our fractions, rather than substantially changing them. This statement is particularly true for the highly enriched genes in the HS fraction. However, we observed only ∼30% of overlap between moderately enriched genes in the wild type and *chchd4a*^-/-^ HS and MB fractions, respectively. At the same time the *chchd4a* loss of function triggers exclusive enrichment of 67 and 188 highly genes in the MB and HS fractions, respectively. As indicated by the KEGG pathway enrichment, *chchd4a*^-/-^ seems to affect the energy metabolism (Figure S8A). Genes detected in the HS fraction of *chchd4a*^-/-^ mutants were directly involved in the oxidative phosphorylation and citrate cycle. At the same time, we observed a handful non-mitochondrial genes (Figure S8B) and no mitochondrial genes (Figure S8C) switching the enrichment between wild type and *chchd4a*^-/-^ mutant MB and HS fractions.

Analysis of the 5’UTR regions of genes exclusively enriched in the HS upon *chchd4a* loss revealed a sequence element that might have increased their presence there. The Translation Initiator of Short 5’ UTR (TISU) element^77,78^ is known to regulate translation of mitochondrial genes, as it confers resistance to translation inhibition in response to energy stress^79,80^. Therefore, we suspect that TISU elements may play a regulatory role in the translation of mitochondrial genes upon *chchd4a* loss of function. As expected, we found the TISU element (SAAS**AUG**GCGGC) highly enriched among the 5’UTR sequences of genes exclusively enriched in the HS fraction (E-value = 2.1 x 10^-14^) of the *chchd4a*^-/-^ mutants (Figure 6F). We also noticed that in *chchd4a^-/-^* mutants, the 5’UTRs of mitochondrial transcripts exclusively enriched in the HS were much shorter compared to those enriched in MB fraction (median = 106 nt vs. median = 166 nt) (Figure 6G). Interestingly, transcripts exclusively enriched in the MB fraction upon *chchd4a^-/-^* mutation had almost 3 times longer CDS sequences compared to transcripts enriched in the HS (Figure S6E). We did not detect any specific sequence motif enriched among the 5’UTRs of genes exclusively enriched in the MB fraction.

## Discussion

Here, we present the genome-wide analysis, which provides new insights into subcellular localization of nuclear-encoded mitochondrial gene products in zebrafish. In this study, we developed a biochemical fractionation method that allows to obtain the membrane-bound fraction containing intact mitochondria and ER with ribosomes on their surface. We showed that this strategy enables to accurately analyze mitochondrial transcripts that are likely to be translated by the free cytosolic and membrane-associated ribosomes. By biochemically fractionating zebrafish 5dpf samples and measuring the RNA abundances in each fraction, we captured the subcellular landscape of nuclear-encoded mitochondrial mRNAs. We also detected various sequence features that may determine the subcellular localization of different mitochondrial transcripts. Moreover, this study describes consequences of proteostatic stress response triggered by the disorders in the mitochondrial protein import for the localization of nuclear-encoded mitochondrial transcripts.

Many studies reported the presence of mRNAs on the mitochondrial surface in yeast^9–14,20,23–25^ and human cell lines^26–28^, but so far no such analysis has been performed *in vivo* in higher eukaryotes. Here, we investigated the population of mRNAs encoding mitochondrial proteins in the mitochondrial vicinity *in vivo* using zebrafish. Our results indicate that the vast majority of mitochondrial transcripts were likely to be translated by the free cytosolic polysomes, whereas just a small fraction of them was detected in the membrane-bound fraction. Interestingly, compared to yeast^7–9^, the population of mitochondrial transcripts located in the proximity of the mitochondrial surface in zebrafish was much more modest. We could not confirm the presence in the membrane-bound fraction for the majority of mitochondria enriched transcripts that were reported for yeast (Figure 2E and Figure S3). On the other hand, the vast majority of mRNAs reported to be located in the proximity of mitochondrial surface in human cell lines^28^ were also enriched in our MB fraction. Our data suggest that, in vertebrates, mitochondrial membrane associated transcripts mainly consisted of mRNAs with specific properties, including long coding sequences and long 3’UTRs (Figure 3D). In addition, we identified three new sequence motifs that are enriched in 3’UTR regions of membrane associated mRNAs encoding mitochondrial proteins in wild type and *chchd4a^-^/^-^* samples, respectively. This opens new questions on the factors and exact mechanism involved in mRNA targeting to mitochondrial membrane.

Our results suggest that the post-translational pathway is likely the main route for mitochondrial proteins, while the co-translational import is rather a specialized path that restricts itself to transporting mainly large proteins with particular properties for which the import of nascent protein chains could be inefficient or problematic for the cell. The observation that genes encoding IMS proteins were mainly enriched in the HS fraction suggests that IMS proteins are likely to be translated by free cytosolic polysomes and imported into IMS after completing their synthesis in the cytosol. This is in line with expectations, as the Mia40-mediated import of IMS proteins import involves oxidation-mediated compaction and requires those proteins to be of very small size^81^. On the other hand, we noticed that large proteins with highly hydrophobic transmembrane domains, including ABC transporters and solute carriers, were enriched in the membrane-bound fraction. Coupling translation with protein import is expected to protect the cell from the toxic effects of protein accumulation or erroneous incorporation of those proteins into other membranes. Therefore, this suggests that the protein properties encoded in the primary transcript sequence might determine the choice of import route already at the mRNA stage. Another important feature of transcripts associated with membranes is their high evolutionary conservation, while transcripts translated by the free cytosolic polysomes are much more rapidly evolving (Figure 3E). This finding is in line with previous reports and might be one of the strategies ensuring the fidelity during mitochondrial protein import by enabling only the co-translational uptake of stable, core mitochondrial proteins^3^.

Our RNA-seq based findings are supported by the experimental validation using both RT-PCR and fluorescent *in-situ* hybridization by RNAscope technique^67,68^ (Figure 4). Interestingly, we noticed the enrichment of *slc16a1a* gene in the MB fraction. The RNA scope together with RT-PCR results confirmed the mitochondrial localization of this gene product. The SLC16A1 (MCT1) is a proton-dependent monocarboxylate transporter, involved in transportation of lactate and pyruvate, across the plasma membrane. Although the SLC16A1 was initially annotated as a mitochondrial protein according to the MitoCarta 2.0 gene set^53^, as a plasma mebrane protein, it was excluded from its updated version of the MitoCarta 3.0^82^. Despite the absence of this protein, in the current MitoCarta3.0 database, there is evidence for localization of SLC16A1 into mitochondria. For example, Brooks’s lab showed that in rat skeletal muscle^71^ and neurons of rat brain regions^83^ SLC16A1 can be expressed both in plasma and mitochondrial membranes. This study indicates that SLC16A1 can facilitate both intracellular and cell-cell lactate shuttles and together with mitochondrial lactate dehydrogenase (LDH) can be involved in lactate oxidation. It is worth to mention that above results solely refer to the SLC16A1 localization at the protein level, whereas in our study we study RNA localization. Interestingly, APEX-seq data revealed that SLC16A1 mRNA localize to specific subcellular compartments, mainly Endoplasmic Reticulum Membrane (ERM) and Outer Mitochondrial Membrane^28^. Dual subcellular localization of *SLC16A1* transcripts can be explained by the large interactions and physical contacts between ER and mitochondria in the cell. Nevertheless, the full understanding of mechanisms driving co-localization of SLC16A1 transcript with the mitochondrial membrane requires further investigation.

Biogenesis of fully functional mitochondria depends on a balance between synthesis and degradation of many cellular proteins^29^. To understand whether stress conditions can increase the interaction between two types of import, we investigated the dynamics of subcellular localization of mRNAs encoding mitochondrial proteins using *chchd4a^-^/^-^* zebrafish mutant line^39^. Surprisingly, disorders in mitochondrial IMS protein import triggered by the disruption of MIA pathway further restricted the membrane associated population of mRNAs to mainly transcripts with the longest coding (Figure S6E) and the most evolutionarily conserved sequences (Figure S6F). At the same time, we observed the reduction of the length of coding sequences for transcripts enriched in the HS fraction upon the loss of *chchd4a* function. The MB and HS enriched transcripts populations differed with respect to their CDS length. The CDSs of transcripts enriched in the MB were 3 times longer as for the ones enriched in HS fraction (Figure S6E). We also noticed that *chchd4a* loss of function additionally increased the HS enrichment of MIA pathway targets that are known to be small-sized proteins (Figure 5C). Interestingly, it has been recently described that mRNA association with mitochondria differs between fermentative and respiratory conditions in yeast^25^. A switch to the respiratory conditions enhances the localization of certain nuclear-encoded mRNAs to the surface of mitochondria and helps to increase the protein synthesis thorough mRNA localization.

Our results indicate that 60% (48) and almost 80% (193) genes found in MB and HS fractions, respectively are present in both wild type and *chchd4a^-^/^-^* samples. This suggests that the stress response triggered by the disruption of mitochondrial import in *chchd4a^-^/^-^* mutants further enhanced the presence of specific transcript species, rather than changing it completely. This statement is particularly true for HS fraction. Detailed analysis revealed that many transcripts exclusively enriched in *chchd4a^-^/^-^* mutants were involved in energy metabolism (Figure S8A) and their increased abundance in the HS fraction coincide with the presence of the Translation Initiator of Short 5’ UTR^77,78^ (TISU) element in the 5’UTR regions of their transcripts. TISU confers resistance to translation inhibition in response to energy stress and is known to regulate the translation of mitochondrial genes^79,80^. Moreover, a recent study investigating the cellular temporal proteostatic stress response in human cell lines, reported proteins localized to mitochondria as the only category of genes significantly upregulated in translation in response to ER stress^84^. The authors showed that the examination of the 5’ UTRs revealed the presence of TISU elements in Complex I-IV mitochondrial genes up-regulated by the ER stress response.

The *chchd4a^-^/^-^* mutation triggers similar transcriptomic changes as the tunicamycin induced ER stress^84^. In both cases, cells struggle with the accumulation of misfolded proteins that in the case of the ER stress is repressing translation of TCA cycle genes. We could not confirm reduced levels of TCA cycle enzymes at the protein level using our mass spectroscopy data from previous study^85^. Moreover, we observed slightly increased (log_2_FC < 0.5) expression of some TCA cycle genes (data not shown). At the same time, we noticed increased activity of one-carbon metabolism (Figure 5B-C) and serine biosynthesis, which can suggest that cells are trying to this alternative route for the NADH production. We also noted that *chchd4a^-^/^-^* mutation enhances the amino acid activation. This is in line with previous findings reporting this upregulation for the ER stress response, so far without any possible explanation^86,87^.

To summarize, this study provides new insights into mitochondrial mRNA localization under physiological and pathological conditions resulting from disruption of mitochondrial import pathway. Contrary to our initial hypothesis and recent reports in yeast^25^, *in vivo* analysis in zebrafish revealed that localized translation are not the predominantly utilized mechanism for the synthesis of mitochondrial proteins.

## Acknowledgements

We thank Drs. M. Macias and T. Węgierski (International Institute of Molecular and Cell Biology in Warsaw) for assistance with microscopy and RNA localization analysis. We show appreciation to Prof. Roderic Guigo (Centre for Genomic Regulation, Barcelona, Catalonia, Spain), Prof. Ulrike Topf and Dr. Michal Turek (Institute of Biochemistry and Biophysics, Polish Academy of Sciences in Warsaw) for fruitful discussions, Mark Diekhans (University of California, Santa Cruz) and Rohit Suratekar (International Institute of Molecular and Cell Biology in Warsaw) for the help with data analysis. We are grateful to members of the Chacinska, Uszczynska-Ratajczak and Winata labs for useful insights. This work was supported by the POLONEZ Fellowship of National Science Centre, Poland, 2016/23/P/NZ3/03730. This project has received funding from the European Union’s Horizon 2020 research and innovation programme under the Marie Skłodowska-Curie grant agreement No 665778 (BUR). This work was supported by EU/FP7: Research Potential FISHMED, grant number 316125.

## Author’s contributions

A.C. and C.L.W. designed and supervised the project; S.S. and B.U.R designed the experiments together with A.C. and C.L.W.; S.S. and M.K. performed the experiments; B.U.R. and M.M. performed bioinformatic analyses; all authors discuss and interpret the data; B.U.R and S.S. wrote the manuscript with the help of A.C. and C.L.W. and contributions from all authors.

## Supplementary figure legends

**Figure S1.**
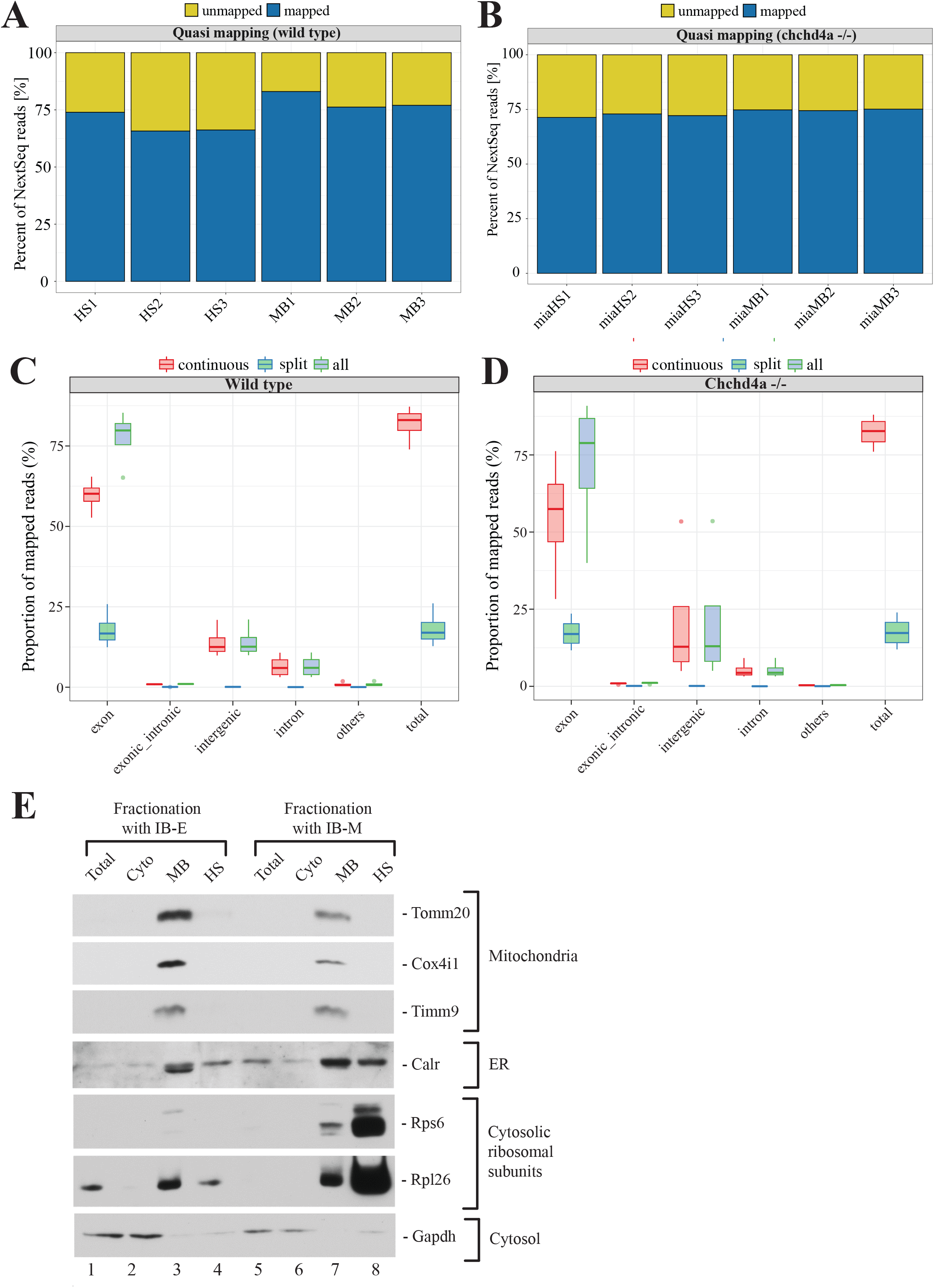
Read-to-transcriptome alignment statistics for wild type (A) and *chchd4a^-^/^-^* (B) samples. Genome coverage for (C) wild type and (D) *chchd4a^-^/^-^* samples. (E) Validation of subcellular fractionation by western blotting. Samples were obtained from 5dpf larvae. IB-E-isolation buffer containing EDTA, IB-M-isolation buffer containing MgCl2 and CHX.

**Figure S2.**
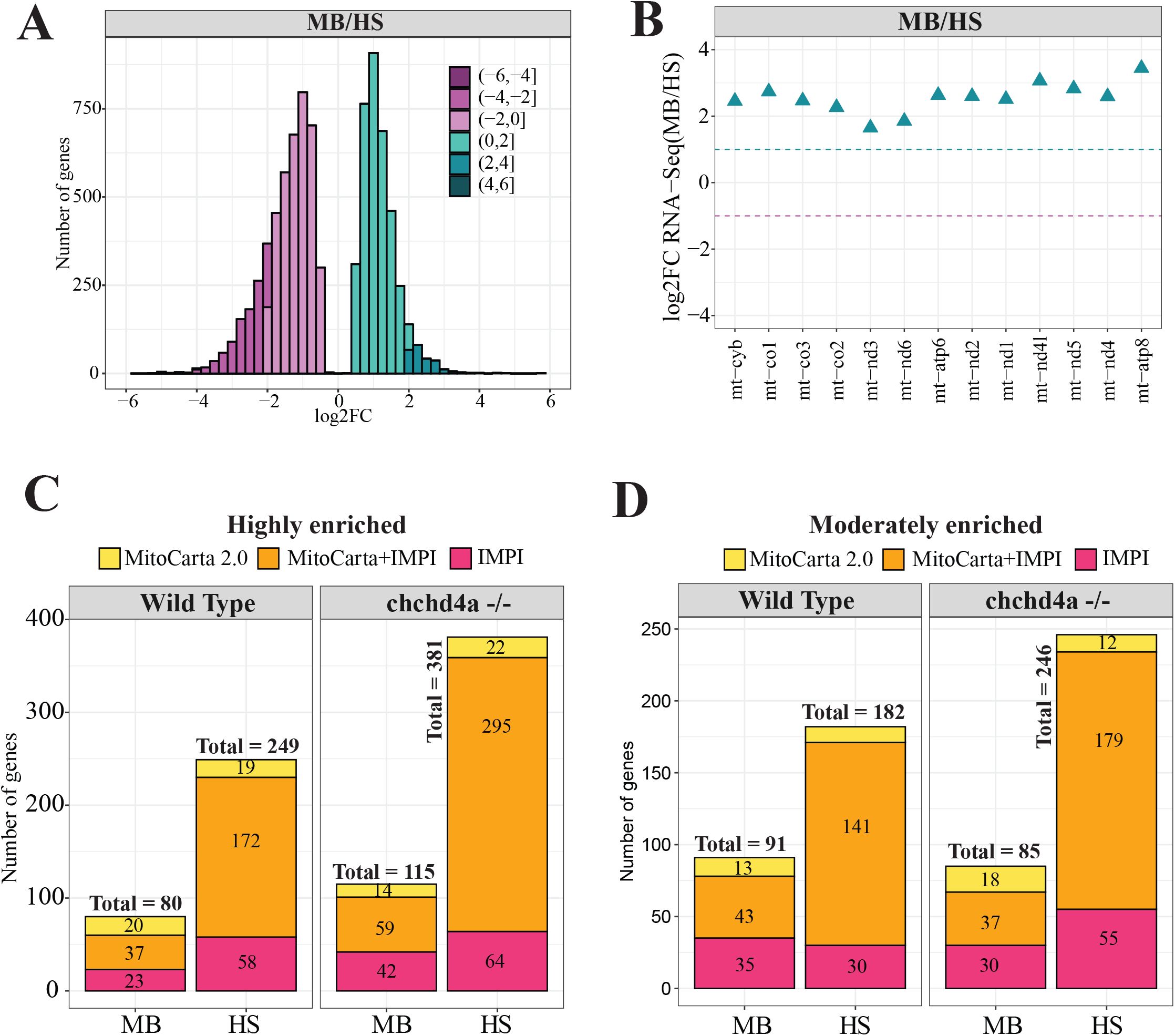
(A) Distribution of log_2_ gene enrichment values for wild type samples. (B) Log_2_ gene enrichment (wild type samples) for genes encoded by the mitochondrial genome. (C) Venn diagrams depicting the overlap between the MitoCarta 2.0 and IMPI gene repositories for MB and HS fractions obtained from wild type and *chchd4a*^-/-^ samples, respectively.

**Figure S3.**
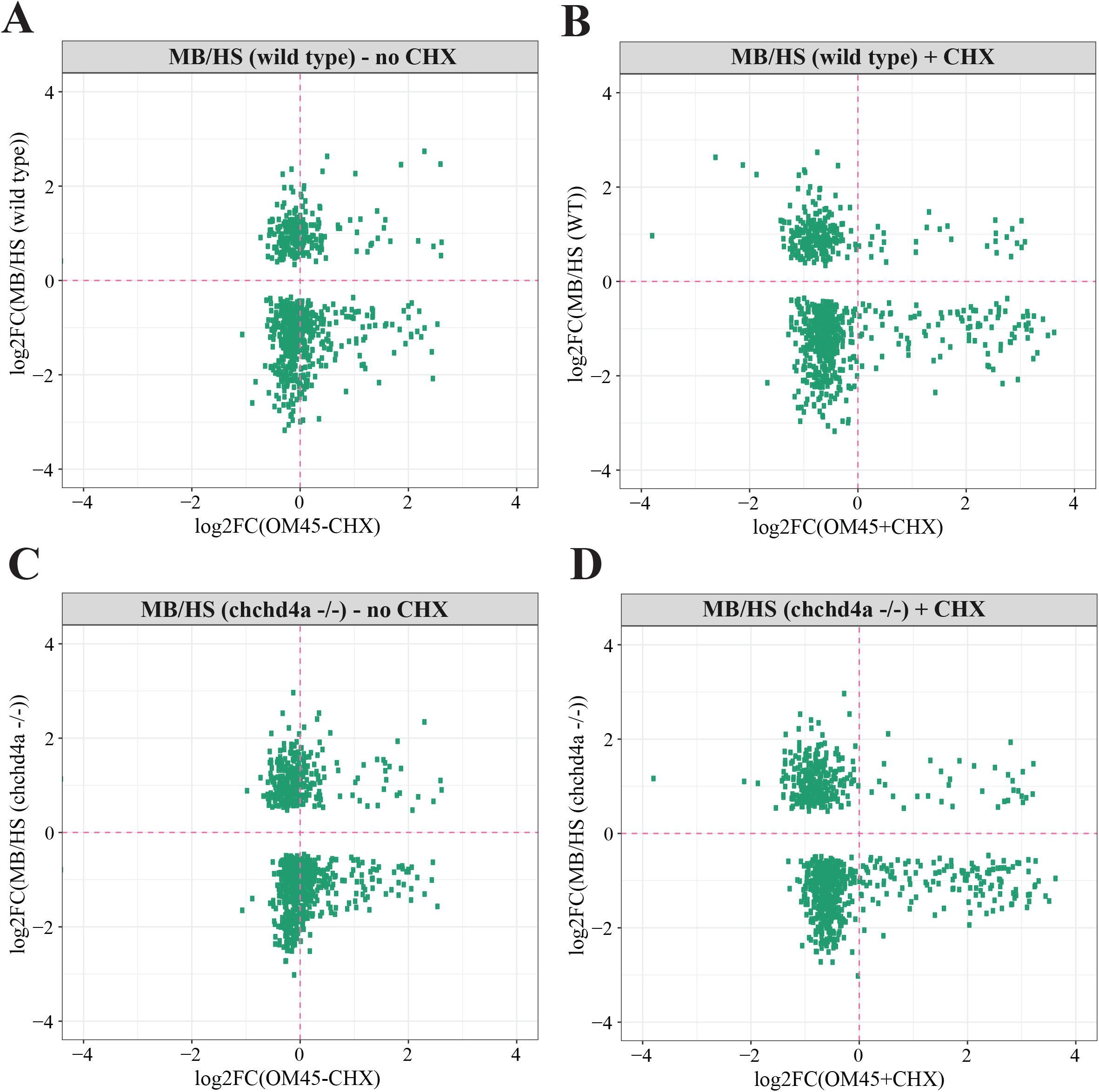
Wild type and *chchd4a*^-/-^ log_2_ gene enrichment values for yeast orthologous genes reported to be enriched at the mitochondrial surface by the proximity specific ribosome profiling with and without CHX^20^.

**Figure S4.**
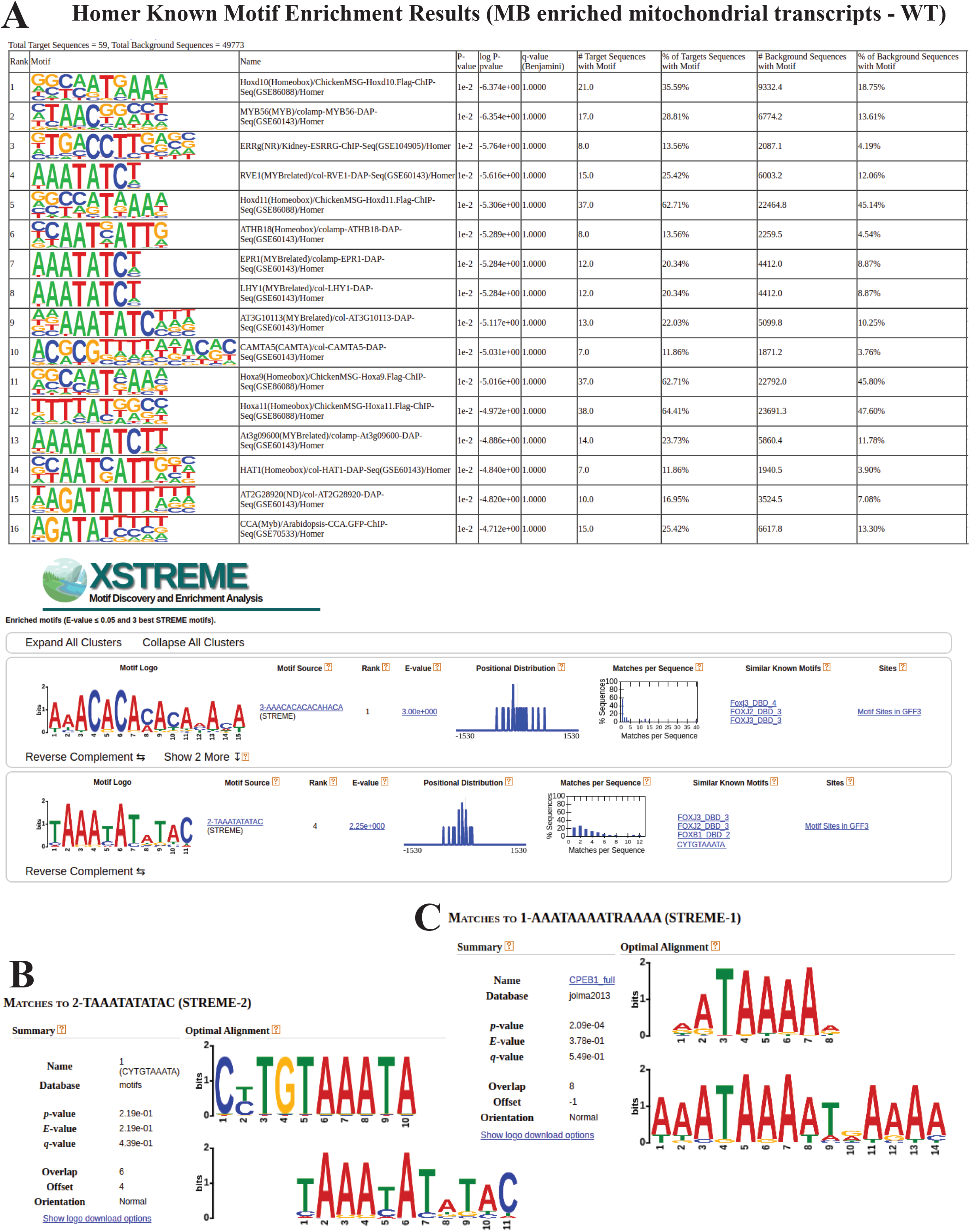
Sequence motifs predicted for mitochondrial genes enriched in the MB fraction (A) by the Homer and STRAME software. (B) Motif comparison with Tomtom software.

**Figure S5.**
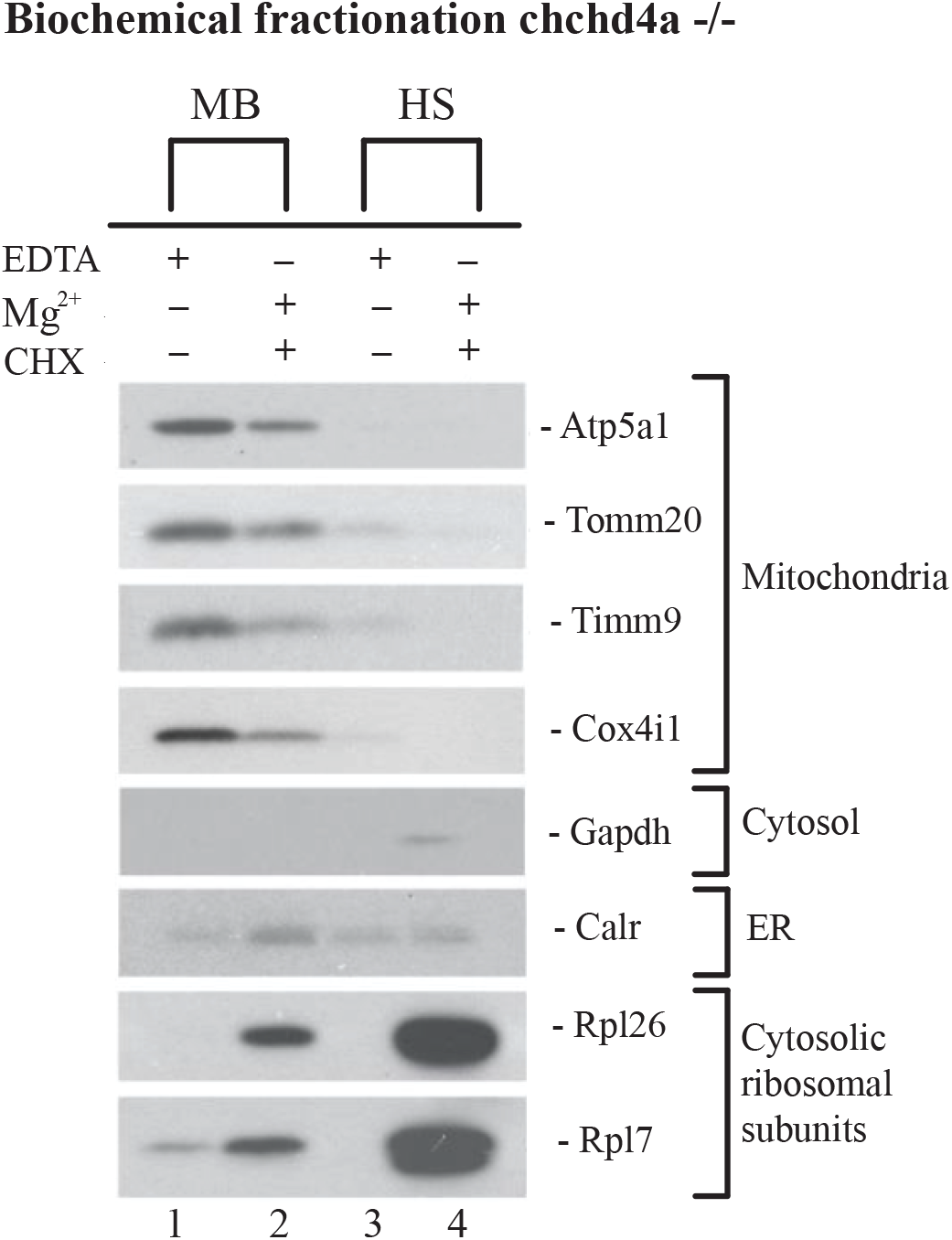
Isolation of membrane-bound and high speed fractions from *chchd4a^-^/^-^* samples.

**Figure S6.**
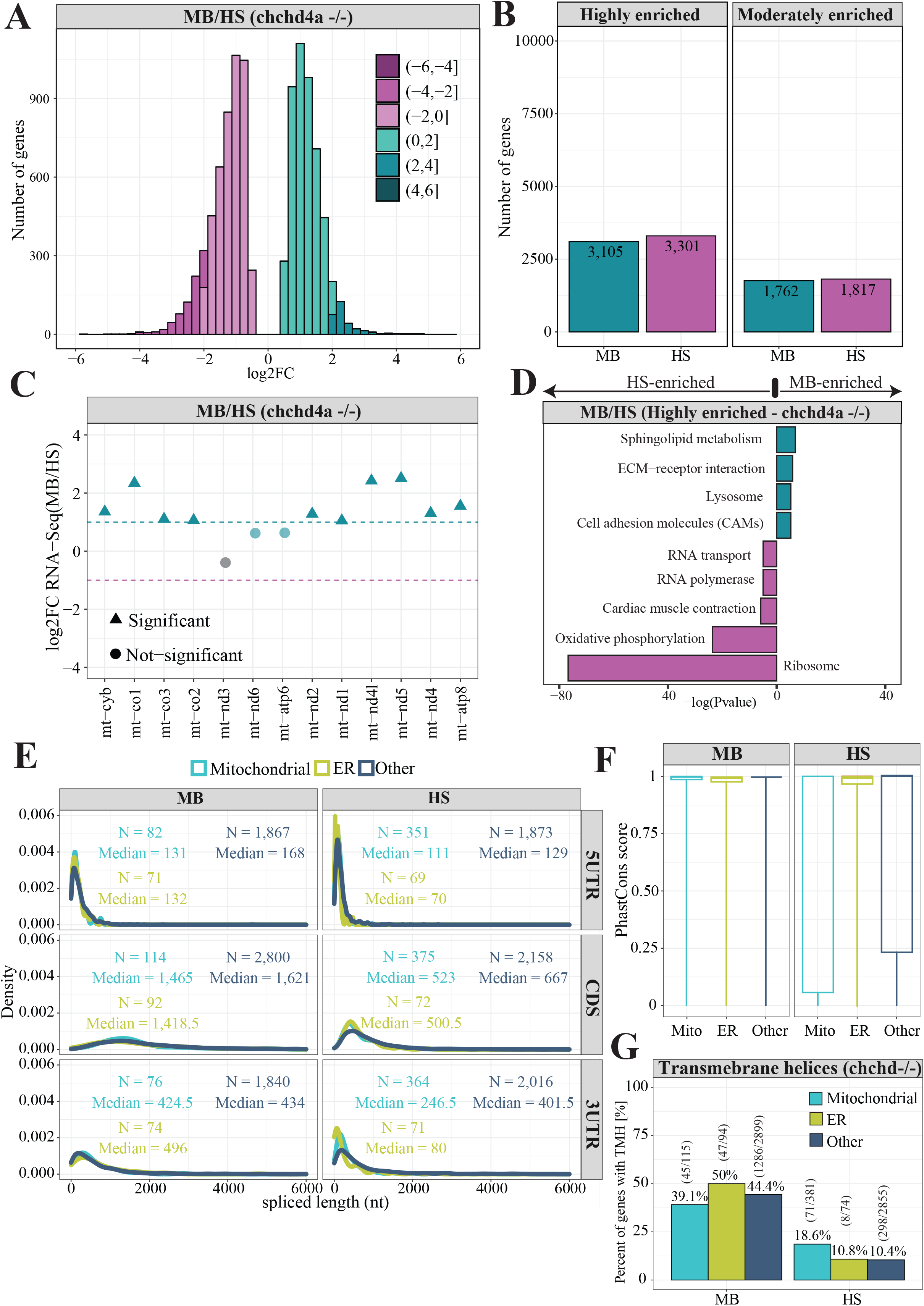
(A) Distribution of log_2_ gene enrichment values *chchd4a^-^/^-^* HS samples (B) Bar plots representing the total number of genes enriched in MB and HS fractions obtained via biochemical fractionation from chchd4a*^-^/^-^* 5 dpf zebrafish larvae. (C) *chchd4a^-^/^-^* log2 gene enrichment values for genes encoded by the mitochondrial genome. (D) KEGG enrichment analysis for genes detected in chchd4a*^-^/^-^* 5 dpf zebrafish samples. (E) Distribution of CDS, 5’ and 3’UTR lengths for transcripts encoding mitochondrial (cyan), ER (green) and other proteins (navy) (F) Boxplots showing PhastCons^48^ conservation tracks for three gene categories. (G) Bar plot representing proportion of transcripts encoding mitochondrial, ER and remaining (other) proteins with transmembrane domains.

**Figure S7.**
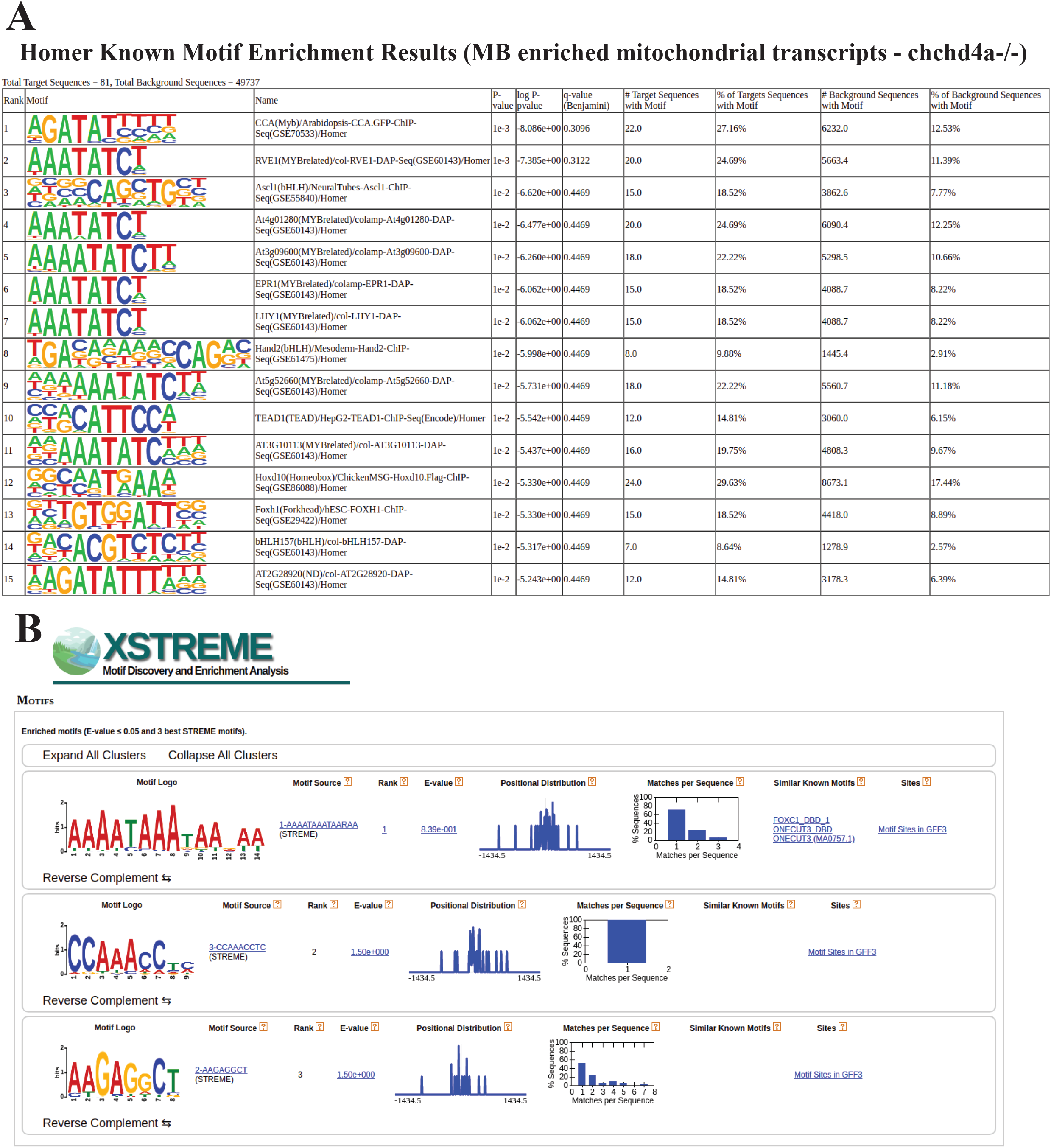
Sequence motifs predicted for mitochondrial genes enriched in the *chchd4a^-^/^-^* MB fraction (A) by the Homer and (B) STRAME software.

**Figure S8.**
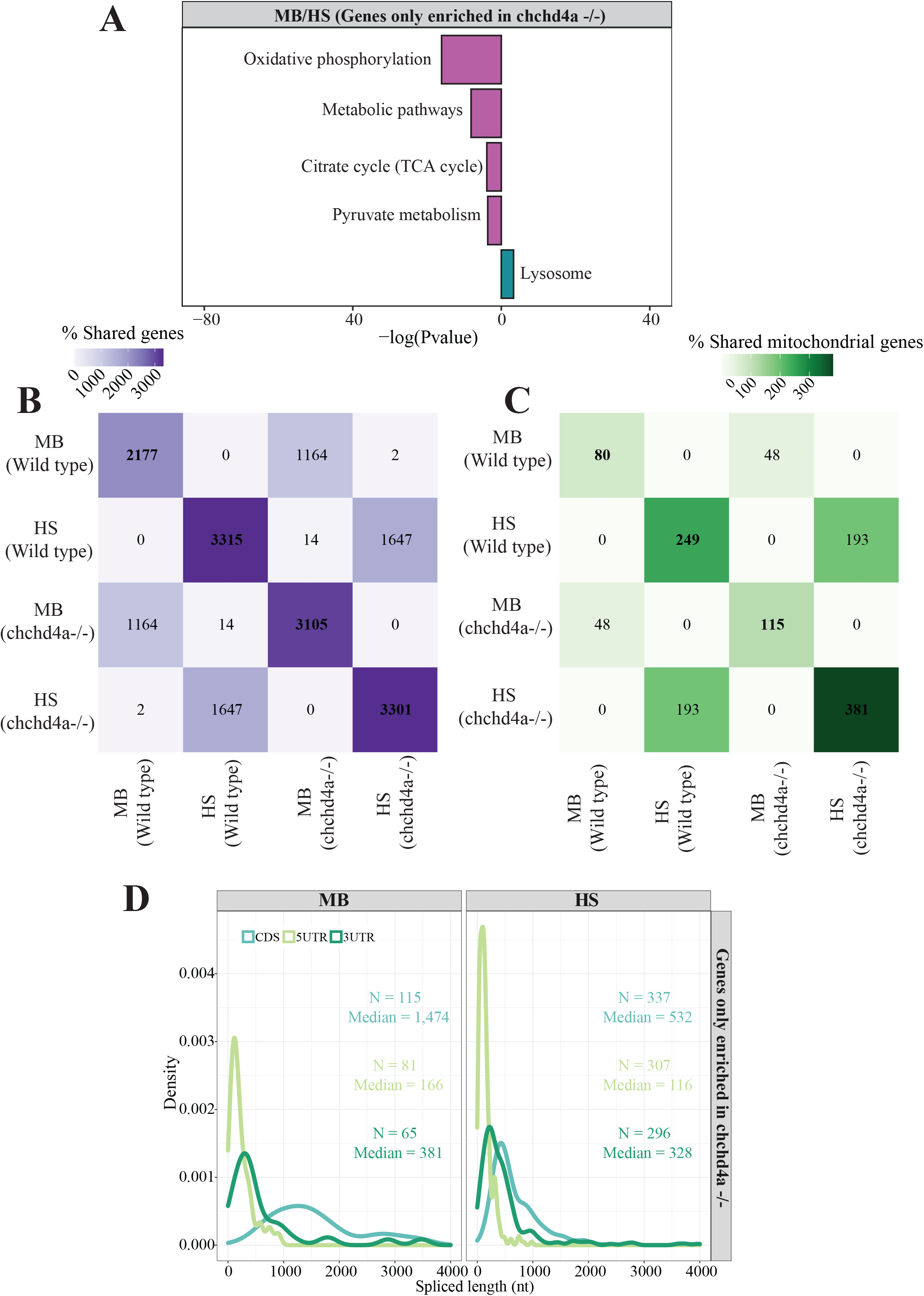
(A) KEGG enrichment analysis for genes exclusively enriched upon chchd4a mutation. Heat maps indicating overlap between all (B) and (C) mitochondrial genes enriched in MB and HS fractions for wild type and *chchd4a^-^/^-^.* (D) Distribution of CDS, 5’ and 3’ UTR lengths for transcripts with TISU element detected in their 5’UTR regions.

## Supplementary tables

**Table S1. List of primers used in this study**

**Table S2. Analysis of RNA sequencing results of wild type and chchd4a^-^/^-^ MB and HS fractions.** Expression matrices for wild type and chchd4a^-^/^-^ samples, highly and moderately enriched genes are shown in separate sheets.

**Table S3. Analysis of RNA sequencing results of *chchd4a*^-^/^-^ mutants and wild-type (WT) controls.**

**Table S4. Human orthologues of zebrafish genes obtained using BioMart data mining tool from Ensembl.**

**Table S5. Zebrafish orthologues of 483 human genes that encode proteins localizing to the ER according to the analysis performed by the Human Protein Atlas^57^.**

## Notes

### Competing Interest Statement

The authors have declared no competing interest.

